# Fibroblast-IL-7 feedback drives PDPN^+^ monocytes to restrain CD4^+^ T cell responses

**DOI:** 10.64898/2026.03.24.713728

**Authors:** Yan Fu, Christophe Toussaint, Ina Sauerland, Elisabeth Reiß, Lars Philipsen, Patricia Gintschel, Laura Knop, Andreas Braun, Bastian Kruse, Leon-Alexander Dewitz, Jonas Amore, Vladyslava Dovhan, Sönke Weinert, Nouria Jantz-Naeem, Iris Baars, Juliane Mohr, Julia Prein, Cora Schwendele, Philipp Bruno, Martin Böttcher, Nicole Gröger, Hannah Van Hove, Melanie Greter, Burkhard Becher, Sascha Kahlfuß, Burkhart Schraven, Robert Geffers, Thomas Tüting, Nicole Joller, Stephan Fricke, Antoine-Emmanuel Saliba, Thomas Schüler, Andreas J. Müller

## Abstract

During immune responses, pro-and anti-inflammatory mechanisms must be balanced to ensure pathogen clearance while limiting tissue damage. Monocyte-derived cells contribute to both processes, yet the underlying regulatory circuits remain incompletely defined. Here, we show that a subset of PDPN^+^IL-7R^+^ monocyte-derived cells that impair effector CD4^⁺^ T cell–mediated control of intracellular pathogens, and thus perpetuate the infection. Fibroblast-derived IL-7 drives this immunosuppressive program, which is up-regulated in response to IFNγ. We thus uncover a cytokine-dependent feedback circuit in which elevated IFNγ induces IL-7 production by fibroblasts, licensing immunosuppressive monocyte-derived cells that restrain CD4⁺ T cell responses. This mechanism links excessive inflammation to immune suppression at the expense of pathogen control. Targeting this feedback loop may enable therapeutic strategies that enhance antimicrobial immunity while preserving tissue integrity.

## Introduction

Inflammation induces the recruitment of large numbers of monocyte-derived myeloid cells which phagocytose invading pathogens, deliver antimicrobial effector molecules and thus limit pathogen load. However, such cells also exert regulatory functions required for the resolution of inflammation and tissue repair (Guilliams et al., 2018). This is of critical importance for host survival, since unrestrained immune responses endanger tissue integrity and organ function (Ortega-Gomez et al., 2013; Postat et al., 2018; Sugimoto et al., 2016). Intracellular pathogens infecting monocyte-derived cells can interfere with anti-microbial effector functions of their host, thereby establishing long-term persistence and chronic disease (Carvalho et al., 2023; Collins-McMillen et al., 2017; Drewry et al., 2019; Mariotti et al., 2004; Stapels et al., 2018). Furthermore, monocyte-derived cells can acquire an immunosuppressive phenotype that counter-regulates effector functions of other immune cells, or shifts immune responses towards a less inflammatory phenotype (Gundra et al., 2017; Zhang et al., 2022). Elucidating and manipulating these diverse functions holds the potential to improve treatment for infectious diseases (Channappanavar et al., 2016; Dorhoi et al., 2014; Wallis et al., 2023) and cancer (Kruse et al., 2023; Mantovani et al., 2022) but the mechanisms by which such functions are induced and deployed remain incompletely understood.

Upon arrival in the tissue, monocytes adapt their functional properties in response to complex environmental cues, including inflammation-and pathogen-derived signals. This context-dependent adaptation is crucial for a balanced response ensuring pathogen elimination and tissue protection at the same time (Veglia et al., 2021). Thus, within a population of activated monocyte-derived cells, a mixture of pathways triggering inflammatory, antimicrobial, and tissue remodeling functions must be activated (Sanin et al., 2022). Single-cell RNA sequencing (scRNAseq) has been instrumental in dissecting monocytes and their derivatives into phenotypically and functionally distinct subpopulations during infection by intracellular bacterial (Bryson et al., 2019; Hoffman et al., 2021), viral (Baasch et al., 2021) or parasitic (Patir et al., 2020) pathogens, as well as in the tumor microenvironment (Cheng et al., 2021; Mulder et al., 2021). Some of these subpopulations have the capacity to critically divert the course of the disease, even if present only at a small number compared to the total monocyte population (Hoffmann et al., 2022; Zhang et al., 2022). In the course of an infection, the level of complexity is further increased by a significant diversity of virulence, growth rate and invasiveness of individual pathogens, thereby additionally modulating the functional properties of the infected cell type (Avraham et al., 2015; Pisu et al., 2021; Saliba et al., 2016).

To address the differential monocyte-derived cell functions in the context of the diverse pathogen states, we employed *Leishmania major* (*L. major*) infection as a physiological model for infection-induced recruitment and activation of monocytes (Heyde et al., 2018; Romano et al., 2017). In this model system, monocyte-derived cells represent the preferential host cell type for intracellular pathogen proliferation and, at the same time, are critical for controlling the parasite through expression of the inducible nitric oxide synthase (iNOS) in response to T cell-derived Interferon-γ (IFNγ) (De Trez et al., 2009; Leon et al., 2007; Muller et al., 2012; Olekhnovitch et al., 2014). Permanent monocyte recruitment to and maturation at the site of infection results in a large phenotypic and functional diversity. Importantly, the proliferation state of the pathogen correlates with the phenotype and function of the infected monocyte-derived cells (Formaglio et al., 2021; Heyde et al., 2018; Mandell and Beverley, 2017). Also, the composition and polarization of these cell populations has been shown to critically impact on the deployment of the anti-*Leishmania* response, e.g. by entertaining type II immune cell circuitries (Lee et al., 2023), or by modulating the reservoir of monocyte-derived cells permissive for pathogen proliferation (Carneiro et al., 2020). However, the cellular and molecular interactions which enable monocyte-derived populations to impact prolongation versus control of the infection remain to be fully elucidated.

Here, we combined an *in vivo* pathogen proliferation reporter system with scRNAseq to characterize the function of distinct monocyte-derived subpopulations in the context of their interactions with intracellular pathogens. With the help of this approach, we identified a subset of Podoplanin (PDPN)-and Interleukin-7 receptor (IL-7R) expressing cells which are derived from monocytes and preferentially harbor low-proliferating *L. major*. This PDPN^+^IL-7R^+^ population exhibited a hypo-inflammatory phenotype and suppressed effector functions of *L. major*-specific CD4^+^ T cells in response to fibroblast-derived IL-7. IL-7 production was dependent on IFNγ release in the course of the adaptive immune response against *L. major*. Interference with IL-7R signaling or depletion of these cells, as well as *Il7* gene inactivation in fibroblasts, resulted in improved effector T cell responses and subsequent pathogen clearance.

In summary, we provide evidence for an as-yet unknown feedback loop which dampens T cell responses as a result of the inflammation-associated up-regulation of fibroblast-derived IL-7, and the subsequent immunosuppressive activity of IL-7R^+^ monocyte-derived cells. This mechanism can favour the pathogen to establish persistence and could therefore represent a new target for immunotherapeutic approaches.

## Results

### PDPN^+^IL-7R^+^ phagocytes infected with low proliferating *L. major* exhibit a hypo-inflammatory phenotype

During local immune responses, tissue-and inflammation-derived signals modulate the phenotype and function of recruited immune cells (Romano et al., 2017; Saliba et al., 2016; Szabo et al., 2021; Yamada et al., 2020). However, dissecting these environmental cues has remained challenging. We and others have shown previously that intracellular *L. major* proliferation differs considerably between infected monocyte-derived cells recruited to the infection site (Heyde et al., 2018; Mandell and Beverley, 2017). We therefore utilized pathogen proliferation rate as a proxy for host-pathogen interaction cues which define distinct monocyte-derived cell functions *in vivo*. For this, we employed an *in vivo* proliferation assay, which relies on genetically modified *L. major* parasites expressing the photoconvertible green fluorescence protein mKikume (*Lm*^SWITCH^). Upon illumination, mKikume molecules convert from green to red. Consequently, red molecules are preserved in low-proliferating parasites (*Lm*^lo^), but are diluted by newly synthesized green molecules in high-proliferating *L. major* (*Lm*^hi^) (Fig. 1A) (Formaglio et al., 2021).

**Figure 1.**
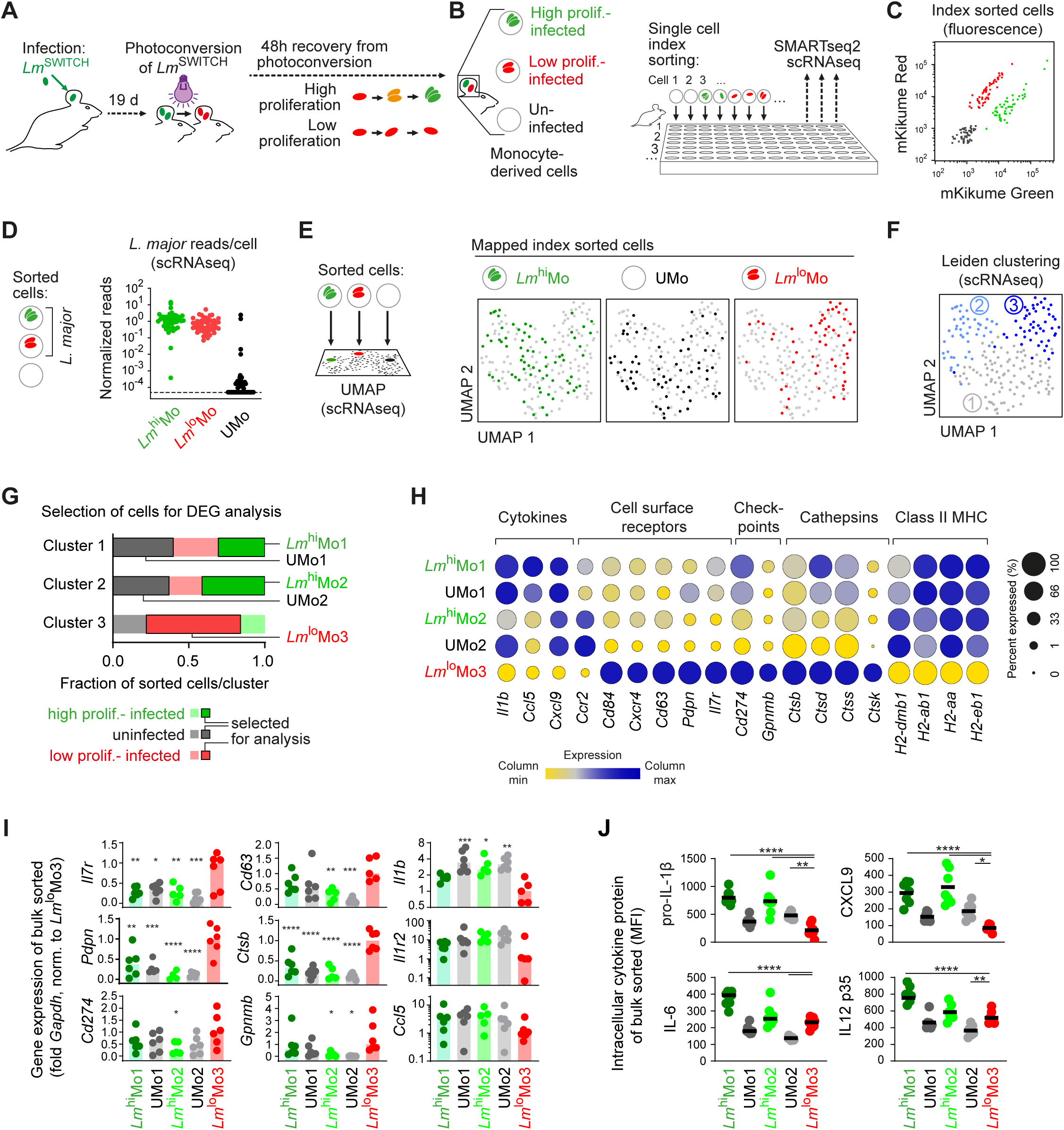
PDPN^+^IL-7R^+^ phagocytes infected with low proliferating *L. major* exhibit a hypo-inflammatory phenotype. **A.** Experimental setup for *in vivo L. major* proliferation measurement using *Lm*^SWITCH^. **B.** Experimental setup for index single-cell sorting of CD11c^int-hi^ and Ly6c^+^ monocytes from 10 individual mouse ears infected with proliferation reporter parasites at 3 weeks post-infection (p.i.). **C.** Index mKikume red and green reporter fluorescence values. **D.** *L. major* reads detected in the sorted monocytes infected with high (green) or low (red) proliferating pathogen, or uninfected (black). **E.** UMAP representation and mapping of cells sorted as high-proliferator infected (green), uninfected (black) and low-proliferator infected (red). **F.** Leiden clustering of the scRNAseq data on the UMAP representation. **G.** Fraction of monocytes sorted as infected with *Lm*^hi^ (green), uninfected (black) or infected with *Lm*^lo^ (red) among the three clusters identified by the Leiden algorithm. The five highlighted most prominent populations (*Lm*^hi^Mo1, UMo1, Cluster 1; *Lm*^hi^Mo2, Umo2, Cluster 2; *Lm*^lo^Mo3, Cluster 3) were selected for differentially expressed gene (DEG) analysis. **H.** Normalized expression (color-code) and cell fraction per population with detected expression (circle size) for selected DEG differentially expressed in *Lm*^lo^Mo3 versus all other populations. **I.** Quantitative PCR (qPCR) analysis of selected genes identified by scRNAseq. Expression levels were determined for replicates of 50-100 cells sorted according to the gating strategy outlined in fig. S1K. Expression levels relative to *Gapdh* and normalized to *Lm*^lo^Mo3 are shown. Each symbol represents 50-100 cell replicates; data was generated from two independent experiments. Horizontal lines denote the median. ****, p<0.0001;***, p<0.001;**, p<0.01;*, p<0.05 compared to the *Lm*^lo^Mo3 population according to one-way ANOVA with Dunnett’s post-test. **J.** Mean fluorescence intensity (MFI) of pro-IL-1β, CXCL9, IL-6, and IL-12 p35 among the *Lm*^hi^Mo1, Umo1, *Lm*^hi^Mo2, Umo2, and *Lm*^lo^Mo3 (according to the gating strategy shown in fig. S1K) from *L. major* infected C56BL/6J mouse ear skin at 3 weeks p.i. Each dot represents one infected ear analyzed, data pooled from two independent experiments. Horizontal lines denote mean. ****, p<0.0001; **, p<0.01; *, p<0.05 according to one-way ANOVA multiple comparison with Dunnett’s post-test.

To characterize the transcriptome of cells which harbour *Lm*^lo^ and *Lm*^hi^, we infected C57BL/6 (B6) mice with 2 x 10^6^ *Lm*^SWITCH^ in the ear. At 19 days post infection (p.i.), the parasites were photoconverted and CD45^+^ leukocytes were MACS purified 2 days later from the infected ear. Subsequently, single-cell sorting of uninfected, *Lm*^lo^-or *Lm*^hi^-containing monocyte-derived cells was performed (Fig. 1B). Both CD11c^int-hi^ and Ly6c^+^ were selected to cover a broad spectrum of recruited monocyte-derived cells (Fig. S1A) (Heyde et al., 2018; Olekhnovitch et al., 2014). *Lm*^lo^-or Lm^hi^-infected (termed *Lm*^lo^Mo or *Lm*^hi^Mo, respectively) as well as uninfected (termed UMo) single monocyte-derived cells were separated using index-sorting into individual wells (Fig. 1C and S1B). ScRNAseq using the SMARTseq2 protocol (Picelli et al., 2013) revealed specifically and consistently high *L. major* transcript counts in cells sorted as infected, but not uninfected (Fig. 1D). In total, 88% of the sorted cells had passed quality control based on mitochondria read counts and percentage of ERCC (Fig. S1C, and see details in Methods).

Mapping of the sequenced cells sorted as *Lm*^hi^Mo (green), UMo (black) and *Lm*^lo^Mo (red) to a batch-corrected Uniform Manifold Approximation and Projection (UMAP) representation (Fig. 1E, S1D, and S1E) revealed that cells infected with *Lm*^lo^ exhibited a unique gene expression profile. In contrast, uninfected cells and cells infected by *Lm*^hi^ were not distinguishable on the UMAP representation. Unsupervised clustering using the Leiden algorithm identified three distinct clusters of cells referred to as Cluster 1, 2 and 3, with Cluster 3 consisting mainly of *Lm*^lo^Mo, (Fig. 1F; Table S1).

Gene ontology (GO) function terms revealed a differential regulation of mainly immune system-related processes for Cluster 3, but not Cluster 1 and 2. In particular, biological process enrichment in Cluster 3 included immunosuppressive signaling pathways, but lowered immune-activating functions and antigen processing as compared to Cluster 1 and 2 (Fig. S1F-G; Table S2-3). Regarding differentially expressed genes (DEG) specific for Cluster 3, we found a strong reduction of the proinflammatory cytokine gene *Il1b*, chemokines and class II MHC-related genes as compared to Cluster 1 and 2. In contrast, cathepsins and immune checkpoint ligands were expressed specifically in Cluster 3, in line with a hypo-inflammatory phenotype suggested by GO terms. Moreover, we found a specific expression of *Pdpn* in Cluster 3, which is reported to be involved in immunomodulatory functions of myeloid cells (Hovd et al., 2024; Rayes et al., 2017). Additionally, *Il7r* was strongly upregulated in Cluster 3. In contrast, *Ccr2* expression was virtually absent in this population (Fig. S1H).

As Cluster 3 consisted mainly of *Lm*^lo^Mo, while *Lm*^hi^Mo and UMo were evenly dispersed across Cluster 1 and 2 (see Fig. 1E), we defined the five most prominent populations for further analysis, i.e. *Lm*^hi^ infected and uninfected cells in Cluster 1 (termed *Lm*^hi^Mo1 and UMo1) and Cluster 2 (termed *Lm*^hi^Mo2 and UMo2), as well as cells infected with *Lm*^lo^ in Cluster 3 (termed *Lm*^lo^Mo3) (Fig. 1G). The gene expression pattern specific for Cluster 3 (see Fig. S1H) was consistent with the comparison of *Lm*^lo^Mo3 versus *Lm*^hi^Mo1, UMo1, *Lm*^hi^Mo2 and UMo2 (Fig. 1H). This notably included the significantly higher expression of *Pdpn* and *Il7r*, suggesting that PDPN^+^IL-7R^+^ *Lm*^lo^Mo3 constitute a hypo-inflammatory cellular subset. These observations were validated by cells sorted according to their differential expression of CD11c (high in Cluster 1), Ly6c (high in cluster 2, intermediate in infected cluster 1) observed during index sorting, and CCR2 (high in cluster 2) observed in scRNAseq (Fig. S1I, and S1J). Quantitative PCR (qPCR) of cells sorted based according to this strategy (Fig. S1K) reproduced the differential gene expression of *Lm*^lo^Mo3 observed in scRNAseq, including the specific expression of *Pdpn* and *Il7r*, and the hypo-inflammatory phenotype indicated by lower expression of *Il1b*, *Il1r2* and *Ccl5* (Fig. S1I). Also, *Lm*^lo^Mo3 exhibited the lowest levels of pro-IL-1β, CXCL9 protein among all five, and the lowest IL-6 and IL-12 p35 protein among the infected cell subsets (Fig. 1J). Irrespective of whether we gated *Lm*^lo^Mo3 cells from the infection gated according to CD11c, Ly6C and CCR2 (see Fig. S1K), or as CD11b^+^Ly6G^-^SiglecF^-^ monocytes gated for PDPN and IL-7R expression, *Lm*^lo^Mo3, PDPN^+^IL-7R^+^ monocytes, respectively, exhibited a low intracellular pathogen proliferation rate (Fig. S1L-N). These data altogether suggested a PDPN^+^IL-7R^+^ phagocyte population which is infected with low proliferating *L. major* and which exhibits a hypo-inflammatory phenotype.

Recently we demonstrated similarities between monocyte-derived cell activation during the immune response against *L. major* and skin melanoma (Kruse et al., 2023). Accordingly, we also detected a monocyte-derived cell population expressing IL-7R and PDPN in skin melanoma at a rate comparable to monocyte-derived cells in the *Leishmania*-infected skin (Fig. S2A-B). Unsupervised Leiden clustering of published single-cell transcriptome data from tumor-associated monocyte-derived cells (Kruse et al., 2023) confirmed the presence of an *Il7r* and *Pdpn-*expressing monocyte-derived population (Figure S2C). This population exhibited a very similar transcriptional pattern as Cluster 3 from *L. major* infected mice (Figure S2D-E, Table S4-5, see also figure S1G-H for comparison), suggesting the occurrence of IL-7R^+^PDPN^+^ hypo-inflammatory monocyte-derived cells beyond infection settings.

These data altogether suggested a PDPN^+^IL-7R^+^ phagocyte population in the inflamed skin which exhibits a hypo-inflammatory phenotype.

### IL-7R^+^PDPN^+^ *Lm*^lo^Mo3 develop from bone-marrow derived CCR2^+^ monocytes and acquire their phenotype in the infected tissue

To further characterize the IL-7R^+^PDPN^+^ *Lm*^lo^Mo3, we analyzed mannose receptor (CD206) and FcγRI (CD64) expression, which distinguished tissue-resident macrophages, dendritic cells as well as different stages of inflammatory monocytes and monocyte-derived macrophages (Carneiro et al., 2020; Lee et al., 2018; Lee et al., 2023) (Fig. 2A and S3A). Infected IL-7R^+^PDPN^+^ phagocytes consisted mainly of Ly6C^int^ monocyte-derived macrophages, but hardly any tissue-resident and monocyte-derived dendritic cells (TR/MoDC), nor tissue-resident macrophages (TRM) (Fig. 2B and S3B). The same was observed when analyzing phagocytes infected with low proliferating *L. major* (Fig. 2B and S3C-D). Thus independently of the gating strategy, IL-7R^+^PDPN^+^ *Lm*^lo^Mo3 cells seemed to consist mainly of Ly6C^int-lo^, monocyte-derived macrophages.

**Figure 2.**
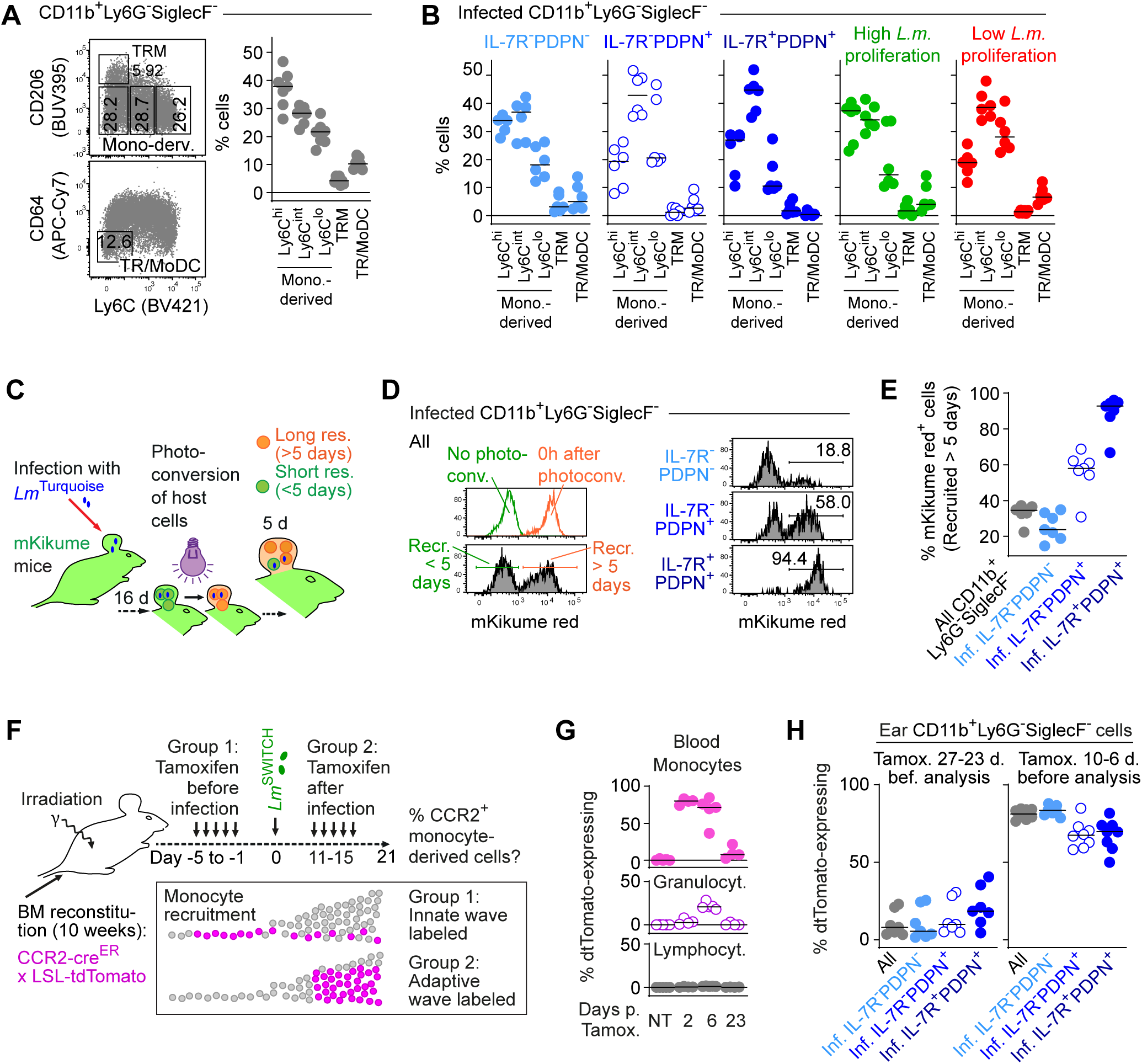
IL-7R^+^PDPN^+^ *Lm*^lo^Mo3 develop from bone-marrow derived CCR2^+^ monocytes and acquire their phenotype in the infected tissue. **A.** Gating to determine tissue-resident and monocyte-derived dendritic cells (TR/MoDC), tissue-resident macrophages (TRM) and different monocyte-derived populations (left) and proportion of these populations among CD11b^+^Ly6G^-^SiglecF^-^ phagocytes (right). **B.** Proportion of the cell populations shown in (A) among *Lm*^hi^Mo1, Umo1, *Lm*^hi^Mo2, Umo2, and *Lm*^lo^Mo3 (H) and among infected CD11b^+^Ly6G^-^SiglecF^-^ cells with differential PDPN and IL-7R expression and different intracellular pathogen proliferation rates. Horizontal lines denote the median. Each dot represents one infected ear analyzed, data pooled from two independent experiments. **C.** Experimental approach to identify newly recruited monocytes by photoconversion of mKikume transgenic mice (left). **D.** Upper left: Infected CD11b^+^Ly6G^-^SiglecF^-^ cells from *Lm*^Turquoise^-infected ear tissue at 3 weeks p.i. from mKikume transgenic mice before (green curve) and 0 h after photoconversion (red curve). Lower left: Identification of short resident mKikume green (recruited <5 days after photoconversion) and long resident mKikume red (recruited >5 days after photoconversion) cells among infected CD11b^+^Ly6G^-^SiglecF^-^ cells. Right: Examples of the proportion long resident cells (mKikume red^+^) among infected CD11b^+^Ly6G^-^SiglecF^-^cells with differential PDPN and IL-7R expression. **E.** Quantification of short and long resident cells among infected CD11b^+^Ly6G^-^SiglecF^-^ cells with differential PDPN and IL-7R expression. Horizontal lines denote the median. Each dot represents one infected analyzed ear, data pooled from two independent experiments. **F.** Experimental approach to demonstrate the origin of *Lm*^lo^Mo3 from bone marrow-derived, CCR2^+^ monocytes recruited during the innate and adaptive phase of the anti-*Leishmania* response. **G.** CCR2-cre^ER^-dependent TdTomato expression in different blood leukocytes cells before, and at day 2, 6 and 23 after the beginning of five daily Tamoxifen injections. Horizontal lines denote the median. Each dot represents one animal, data pooled from two independent experiments. **H.** Fraction of CCR2-cre^ER^-dependent tdTomato expressing cells among PDPN and IL-7R expressing monocyte-derived cells. The cre recombinase activity was induced in CCR2^+^ monocytes for the innate phase of the immune response by injecting Tamoxifen before infection (left), or for the adaptive phase by injection at 11-15 days p.i. (right). Horizontal lines denote the median. Each symbol represents one infection site per condition obtained from two independent experiments.

Having shown a monocyte-derived phenotype, we sought to determine the dynamics of IL-7R^+^PDPN^+^ *Lm*^lo^Mo3 recruitment and turnover. For this, we employed optical time-stamping with mKikume-transgenic mice, in which photoconversion allows the identification of cells recruited before and after a distinct time point (Formaglio et al., 2021; Heyde et al., 2018). The mice were infected with *L. major* expressing blue-fluorescent mTurquoise (*Lm*^Turquoise^), and mKikume-expressing host cells in the ear were photoconverted at 16 dpi, and analyzed by flow cytometry 5 days later (Fig. 2C). This allowed determining IL-7R and PDPN expression in infected cells and to discriminate photoconverted, long resident (recruited > 5 days before analysis) cells from short resident cells which had been recruited after photoconversion (Fig. 2D and S3E). Importantly, PDPN^+^IL-7R^+^ cells were found nearly exclusively in the long resident cell fraction, while all other monocyte-derived cells exhibited a higher turnover rate and were recruited mainly after the photoconversion (Fig. 2D and E).

Because of their long tissue residence time, we aimed to formally demonstrate whether or not *Lm*^lo^Mo3 derive from CCR2^+^ monocytes. For this, we employed lethally irradiated mice reconstituted with Ccr2-creERT2 (Croxford et al., 2015) x Ai14 (Rosa-CAG-LSL-tdTomato-WPRE) BM, in which *Ccr2*-expressing cells permanently express red fluorescent tdTomato upon Tamoxifen treatment. Tamoxifen was applied either before the infection in order to label cells recruited during the innate wave of monocyte recruitment, or from day 11 p.i. in order to label monocytes recruited during the established adaptive immune response (Carneiro et al., 2020) (Fig. 2F). Tamoxifen treatment induced robust labeling of >75% of blood monocytes for several days (Fig. 2G and S3F). In the infected ear, all monocyte-derived populations, including PDPN^+^IL-7R^+^, showed a large proportion of CCR2-dependent tdTomato expressing cells if Tamoxifen was applied during the adaptive wave of monocyte recruitment, but not if the innate wave of monocytes was labeled (Fig. 2H, S3G-H). Therefore, we concluded that despite its low turnover rate as compared to other monocyte-derived cells, the PDPN^+^IL-7R^+^ *Lm*^lo^Mo3 population was derived from CCR2^+^ inflammatory monocytes recruited to the site of infection.

### PDPN^+^IL-7R^+^ *Lm*^lo^Mo3 dampen T cell responses against *L. major* infection *in vivo*

Having shown IL-7R and PDPN expression for monocyte-derived cells in the infected ear, we assessed whether other immune cell types at the infection site, in particular non-myeloid cells, might also include PDPN^+^IL-7R^+^ double-positive cells (Fig. S4A-C). While as expected, IL-7R was expressed on several leukocyte subsets, PDPN expression, and, consequently, PDPN and IL-7R double-expression, was limited to monocyte-derived, in particular infected, cells (Fig. 3A).

**Figure 3.**
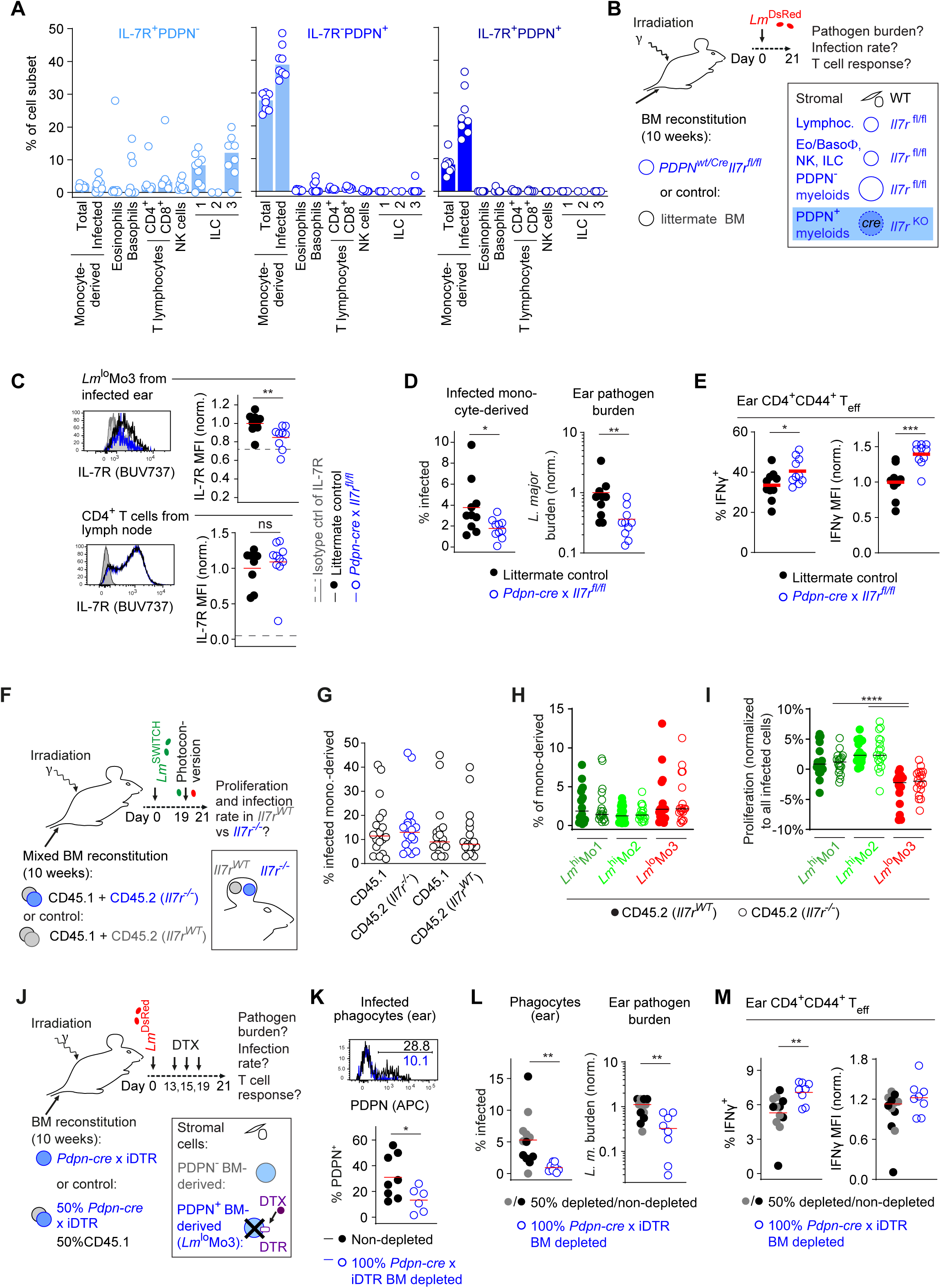
PDPN^+^IL-7R^+^ *Lm*^lo^Mo3 dampen T cell responses against *L. major* infection *in vivo*. **A**. Proportion of PDPN^-^IL-7R^+^ (left), PDPN^+^ IL-7R^-^ (middle) and PDPN^+^IL-7R^+^ (right) fractions among different leukocytes at the infection site at week 3 p.i. Bars denote the median. Each symbol represents one infection site per condition obtained from two independent experiments. **B.** Experimental setup for cell-specific *Il7r*-deficiency in PDPN^+^ leukocytes. **C.** Validation of cell-specific IL-7R deficiency. Upper panels: Example and MFI analysis for IL-7R surface expression on *Lm*^lo^Mo3 (gated according to figure S1K) in the infected ear tissue at 3 weeks p.i. Lower panels: Example and MFI analysis of IL-7R expression on CD4^+^ T cells in the draining lymph node (dLN) at week 3 p.i. Values are normalized to control animals. Each dot represents one infected ear or dLN analyzed. Isotype control-stained samples are shown as a grey histogram and dashed line in the data plot, respectively. Horizontal lines denote mean, data pooled from 2 independent experiments. **, p<0.01; ns, not significant according to unpaired t-test. **D.** *L. major*-infected cell fraction among recruited monocytes in the ear tissue (left) and limiting dilution assay (LDA) of *L. major* tissue burden normalized to control animals (right) at week 3 p.i. of lethally irratiated mice either reconstituted with littermate control (black symbols) or *Pdpn-cre* × *Il7r^fl/fl^*BM (blue symbols). **E.** IFNγ^+^ fraction and IFNγ MFI among effector CD4^+^ T cells within *L. major*-infected ear at week 3 p.i.for mice reconstituted with either littermate control (black symbols) or *Pdpn-cre* × *Il7r^fl/fl^* BM (blue symbols). Each symbol in (D-E) represents one individual mouse ear. Horizontal lines denote the mean. ***, p<0.001; **, p<0.01; *, p<0.05 according to unpaired t-test. Data representative of two independent experiments. **F.** Experimental setup for mixed BM chimera reconstituted with 50% CD45.1 WT and 50% *Il7r^-/-^* (CD45.2) BM. Mice reconstituted with mixtures of WT CD45.1 and WT CD45.2 BM served as controls. **G.** *L. major* infected cell fraction among recruited monocytes in the ear tissue at week 3 p.i., either WT BM chimera compartment (black symbols), or *Il7r^-/-^* BM chimera compartment (blue symbols). **H.** Fraction of *Lm*^hi^Mo1, *Lm*^hi^Mo2, and *Lm*^lo^Mo3 (gated according to figure S1K) among monocyte-derived cells of each BM chimera compartment, either WT BM chimera compartment (closed symbols), or *Il7r^-/-^* BM chimera compartment (open symbols) within *L. major* infected ear tissue at 3 week p.i. Each symbol represents one individual mouse ear, data pooled from two independent experiments. Horizontal lines denote the median. No significance detected according to one-way ANOVA multiple comparison with Dunn’s post-test. **I**. Intracellular pathogen proliferation rates among *Lm*^hi^Mo1, *Lm*^hi^Mo2, and *Lm*^lo^Mo3 (gated according to figure S1K), either WT BM chimera compartment (closed symbols), or *Il7r^-/-^* BM chimera compartment (open symbols). Each symbol represents one individual mouse ear, data pooled from two independent experiments. Horizontal lines denote the median. ****, p<0.0001 according to one-way ANOVA multiple comparison with Dunnett post-test. **J.** Experimental setup for depletion of *Pdpn*-expressing leukocytes of lethally irradiated mice reconstituted with either *Pdpn-cre* × iDTR BM cells, or 50% of *Pdpn-cre* × iDTR and 50% of CD45.1 WT BM cells. **K.** Validation of depletion of PDPN^+^ cells among infected leukocytes. Upper panel: Example of PDPN staining of infected cells from ear tissue at 3 weeks p.i. in depleted (blue curve) or non-depleted control animals (black curve). Lower panel: Percentage of PDPN^+^ cells among infected leukocytes from ear tissue at 3 weeks p.i. in depleted (blue symbols) or non-depleted control animals (black symbols). Horizontal lines denote the mean. Each dot represents one infected ear analyzed. *, p<0.05; ns, not significant according to unpaired t-test. **L.** *L. major* infected phagocyte fraction (left) and *L. major* burden (right, normalized to non-depleted control) in the ear at week 3 p.i. in non-depleted (black symbols) or partially depleted (grey symbols) controls versus 100% *Pdpn*-dependent depleted BM chimeras (blue symbols). **M.** Fraction (left) and MFI (right) of IFNγ^+^ expressing CD4^+^CD44^+^ effector T cells from ear tissue (left) in the ear at week 3 p.i. in non-depleted (black symbols) or partially depleted (grey symbols) controls versus 100% *Pdpn*-dependent depleted BM chimeras (blue symbols). Each symbol in (L) and (M) represents one individual mouse ear, data representative of two independent experiments. Horizontal lines denote the median. **, p<0.01 according to Mann-Whitney test.

Since all IL-7R^+^ monocyte-derived cells co-expressed PDPN, we next employed a *Pdpn-cre* based BM reconstitution approach to specifically abolish IL-7R in these cells (Fig. 3B). For this, we lethally irradiated WT recipient mice, reconstituted them with either *Pdpn-cre* × *Il7r^fl/fl^* or littermate BM, and infected them with *Lm^DsRed^* 10 weeks after reconstitution. Pathogen control and effector T cell activation were analyzed at week 3 p.i. We found that IL-7R was significantly diminished in *Lm*^lo^Mo3, but not in lymph node T cells, confirming monocyte-specific gene deletion of *Il7r* (Fig. 3C and S4D). Strikingly, the *Pdpn-cre* dependent *Il7r* knockout on BM-derived cells significantly reduced *L. major* infected monocytes and pathogen burden (Fig. 3D and S4E). Furthermore, IFNγ production in effector CD4^+^ T cells within the infected ear tissue was elevated (Fig. 3E and S4F), suggesting that disruption of IL-7R signaling in monocyte-derived cells at the site of infection enhances pathogen control by the adaptive immune response. We were able to reproduce the enhanced pathogen control in a different experimental system by reconstituting irradiated recipients with *Rag1^-/-^*× *Il7r^-/-^* BM and adoptive transfer of WT CD4^+^ (Fig. S4G). The resulting animals, in which IL-7R was exclusively present on lymphocytes, also exhibited enhanced control of *L. major,* confirming the role of IL-7R signaling in non-lymphocyte immune cells (Fig. S4H-I).

Enhanced pathogen control in the absence of IL-7R signaling in monocyte-derived cells could be due to cell-intrinsic enhancement of intracellular pathogen control in individual cells devoid of IL-7R, or due to cell-extrinsic effects. In order to address these possibilities, we generated mixed BM chimeras reconstituted with 50% CD45.1 WT and 50% *Il7r^-/-^* (CD45.2) BM. Mice reconstituted with mixtures of WT CD45.1 and WT CD45.2 BM served as controls (Fig. 3F). We found that infection rates (Fig. 3G), the relative abundance, (Fig. 3H), and the parasite proliferation within (Fig. 3I) the different monocyte-derived cell subsets were independent of the cellular *Il7r* genotype in the mixed BM chimeras. Therefore, we concluded that IL-7R signaling in the IL-7R^+^PDPN^+^ *Lm*^lo^Mo3 does not affect their development, abundance, or cell-intrinsic defense function, but dampens pathogen control in a cell-extrinsic manner.

Next, we aimed to assess the effect of the depletion of PDPN^+^IL-7R^+^ *Lm*^lo^Mo3 on the control of *L. major*. For this, we crossed *Pdpn-cre* transgenic mice to iDTR mice expressing the Diphtheria Toxin receptor (DTR) after Cre-mediated elimination of a DNA-STOP cassette (Buch et al., 2005). Using these mice as donors, we created BM chimeras reconstituted with 100% *Pdpn-cre* × iDTR, as well as controls reconstituted with mixed 50% WT (CD45.1) and 50% *Pdpn-cre* × iDTR (CD45.2) BM (Fig. 3J). Diphtheria Toxin (DTX) treatment of mice reconstituted with *Pdpn-cre* × iDTR BM reduced significantly the percentage PDPN^+^ cells among infected phagocytes (Fig. 3K), indicating successful depletion at the site of infection. Importantly, the percentage of infected cells, as well as *L. major* tissue burden was significantly reduced in these animals as compared to non-depleted of only partially depletable BM chimera reconstituted with 50% WT and 50% *Pdpn-cre* × iDTR BM (Fig. 3L and S4J). Of note, this depletion was also associated with an elevated IFNγ production by CD4^+^ effector T cells (Fig. 3M and S4K). These results underlined an immunosuppressive function of PDPN^+^IL-7R^+^ *Lm*^lo^Mo3 in promoting parasite persistence via the inhibition of CD4^+^ T effector functions. Thus, we concluded that IL-7R signaling in *Lm*^lo^Mo3 ultimately resulted in a dampened T cell response against *L. major* and impaired pathogen control.

### IL-7 from IFNγ-induced fibroblasts establishes a feedback loop that limits the effector T cell response in the skin

Stromal cells, particularly fibroblasts, have been shown to produce IL-7 in response to *L. major* infection (Peduto et al., 2009). In line with this, we found that fibroblasts from infected *Pdpn-cre* × *Il7r^fl/fl^* BM reconstituted tissues expressed more MHC-I and PD-L1 than controls, indicating their additional activation due to the enhanced immune response (Figure 4A and S5A). The same was observable in *Pdpn-cre* × iDTR BM chimeric mice in which PDPN^+^ leukocytes were depleted (Fig. 4A and S5B). Of note, we found that *Il7* gene expression was upregulated in the tissue upon *L. major* infection (Fig. 4B and S5C). We therefore sought to elucidate if skin fibroblasts could represent the source of the IL-7 modulating the immunosuppressive function of PDPN^+^IL-7R^+^ *Lm*^lo^Mo3. For this, we employed *Prrx-cre* × *Il7^fl/fl^*mice, which lack *Il7* gene activity in fibroblasts (Knop et al., 2020). Quantitative PCR of stromal cells sorted from the ear confirmed that *Prrx* expression was limited to fibroblasts (Fig. 4C), which were also more abundant in the infected skin than vascular endothelial cells (Fig. 4D and S5D). Additionally, *Il7* expression was significantly reduced in fibroblasts from infected *Prrx-cre* × *Il7^fl/fl^* mouse ear tissue, but not in any other stromal cell population, further confirming a fibroblast-specific *Il7* gene deficiency (Fig. 4E-F). This suggested that *Prrx-cre* × *Il7^fl/fl^* mice indeed lacked *Il7* primarily in fibroblasts in the infected tissue, and that fibroblasts are present in high numbers as compared to vascular endothelial cells as alternative producers of IL-7 (Hara et al., 2012). Importantly, *L. major* clearance was enhanced in *Prrx-cre* × *Il7^fl/fl^* mice (Fig. 4G) and IFNγ production by CD4^+^ T cells was increased (Fig. 4H and S5E). This suggested that fibroblast-derived IL-7 was involved in the suppression of the immune response against *L. major*.

**Figure 4.**
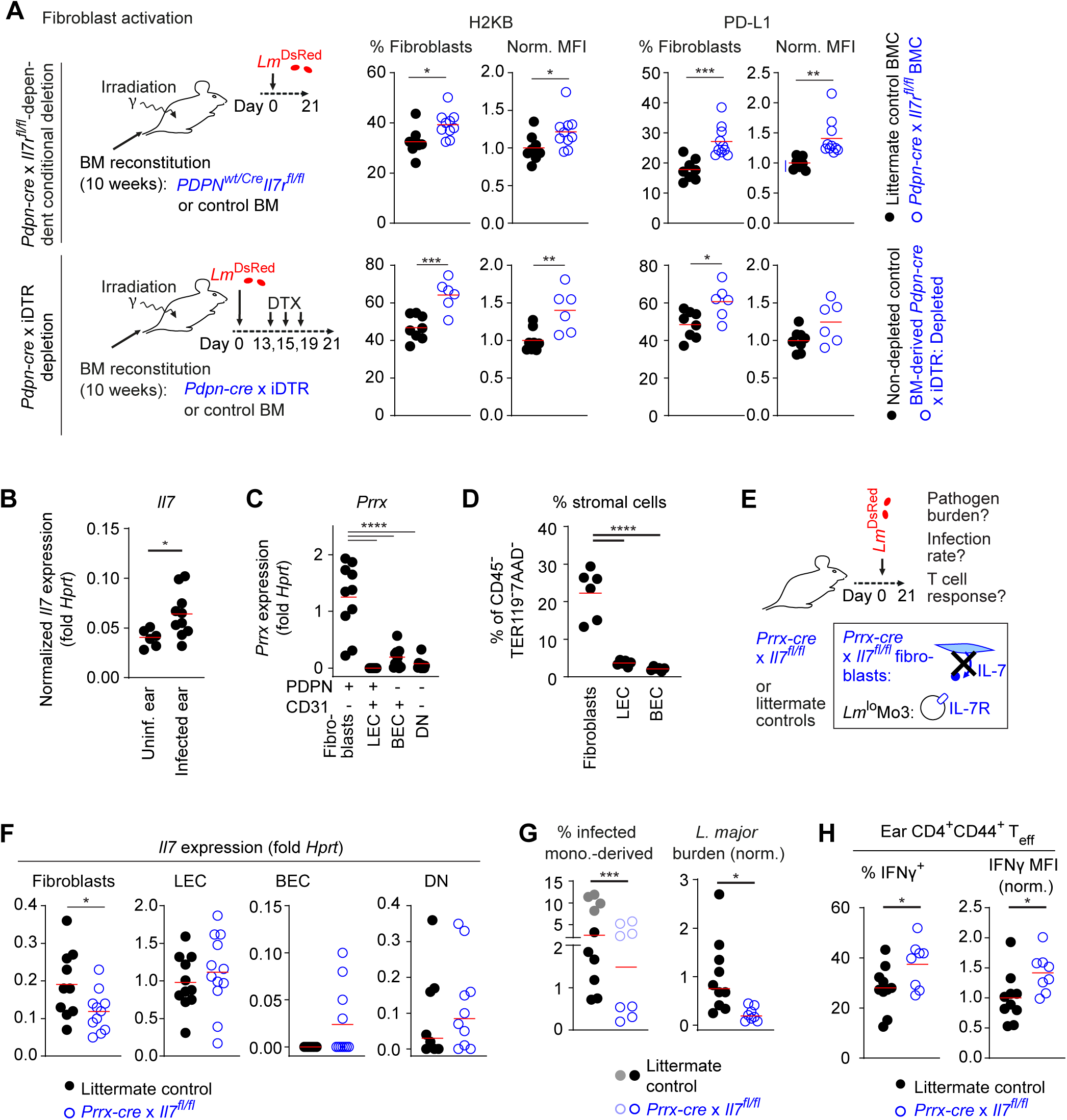
Fibroblast-produced IL-7 contributes to pathogen persistence and suppression of IFNγ production in effector T cells. **A.** Upper panels: H2KB^+^ or PD-L1^+^ fraction and H2KB or PD-L1 MFI of among fibroblasts from mice reconstituted with *Pdpn-cre* × *Il7r^WT/WT^* littermate control BM (black symbols) or *Pdpn-cre* × *Il7r^fl/fl^* BM (blue symbols). Data normalized to mice reconstituted with littermate control BM. Lower panels: H2KB^+^ or PD-L1^+^ fraction and H2KB or PD-L1 MFI of among fibroblasts from non-depleted control (black symbols) or *Pdpn*-dependent 100% depleted (blue symbols) ear infections. Each symbol represents one individual mouse ear. Data normalized to non-depleted control animals. Horizontal lines denote the mean. ***, p<0.001; **, p<0.01; *, p<0.05 according to unpaired t-test. Data shown correspond to the experiments shown in fig. 3C-E, and K, respectively. **B.** Expression of *Il7* within *L. major*-infected (week 3 p.i.) and non-infected ear tissue. Expression as determined by ΔΔCT between *Il7* and *Hprt* qPCR. CT values are corrected according to the mean difference in RNA extracted (see fig. S5C). Each symbol represents one individual mouse ear. Horizontal lines denote the mean. *, p<0.05 according to unpaired t-test. **C.** Expression level of *Prrx* among fibroblasts, LEC, BEC and PDPN^-^CD31^-^non-immune cells within *L. major*-infected ear tissue at week 3 p.i.. Expression as determined by ΔΔCT between *Prrx* and *Hprt* qPCR followed by normalization to PDPN^-^CD31^-^ cells. Each symbol represents one individual mouse ear. Horizontal lines denote the mean. ****, p<0.0001 according to one-way ANOVA multiple comparison with Dunnett’s post-test. Data in (A-C) pooled from two independent experiments. **D.** Proportion of fibroblasts, LEC, and BEC among CD45^-^TER119^-^7AAD^-^ cells within the *L. major* infected ear at week 3 p.i.. Each symbol represents one individual mouse ear. Horizontal lines denote the mean. ****, p<0.0001 according to one-way ANOVA multiple comparison with Dunnett’s post-test. **E.** Experimental setup to test for *Prrx*-expressing fibroblasts as a necessary source of IL-7 to promote *Lm*^lo^Mo3 mediated suppression of CD4^+^ T cell responses in the *L. major* infected ear tissue. **F.** Expression level of *Il7* among fibroblasts, LEC, and BEC within *L. major*-infected ear tissue from littermate control (*Prrx-Cre* × *Il7*^wt/wt^) (black symbols) and *Prrx-cre* × *Il7^fl/fl^* (blue symbols). Expression as determined by ΔCT between *Il7* and *Hprt* qPCR. **G.** Left: the fraction of *L. major* infected cells among recruited monocytes in the ear tissue at week 3 p.i., littermate control (*Prrx-Cre* × *Il7*^wt/wt^) (filled symbols) and *Prrx-cre* × *Il7^fl/fl^*(open symbols). Differently shaded symbols represent two independent experiments. Each symbol represents one individual mouse ear. Horizontal lines denote the median. ***, p<0.001 according to two-way ANOVA comparing genotypes. Right: Limiting dilution assay (LDA) of *L. major* tissue burden in the ear. Data normalized to littermate control animals. **H.** IFNγ^+^ fraction and MFI of IFNγ signal among effector CD4^+^ T cells within *L. major*-infected ear at week 3 p.i. of either littermate control (black symbols) or *Prrx-cre* × *Il7^fl/fl^* animals (blue symbols). Data normalized to littermate control animals. Each symbol in (G-I) represents one individual mouse ear, data are from two independent experiments. Horizontal lines denote the mean. *, p<0.05 according to unpaired t-test.

IFNγ has been reported to induce IL-7 expression in skin fibroblasts (Dai et al., 2021). Therefore, we hypothesized that enhanced IFNγ production by T cells may contribute to a fibroblast-associated feedback loop, limiting the effectiveness of an initially productive T cell response. This may be a result of IL-7 induction by fibroblasts finally leading to immunosuppression of T cells by PDPN^+^IL-7R^+^ *Lm*^lo^Mo3 and subsequent pathogen persistence. In line with this hypothesis, we had shown previously that IFNγ from T cells alone is sufficient for *L. major* containment (Muller et al., 2012), and found T cells represented the main IFNγ producing cell population (Fig. 5A and S6A-B). Also, fibroblasts cultured in the presence of recombinant IFNγ exhibited increased *Il7* cytokine gene expression (Fig. 5B). This suggested that T cell-derived IFNγ could stimulate fibroblasts in the infectious environment to produce IL-7.

**Figure 5.**
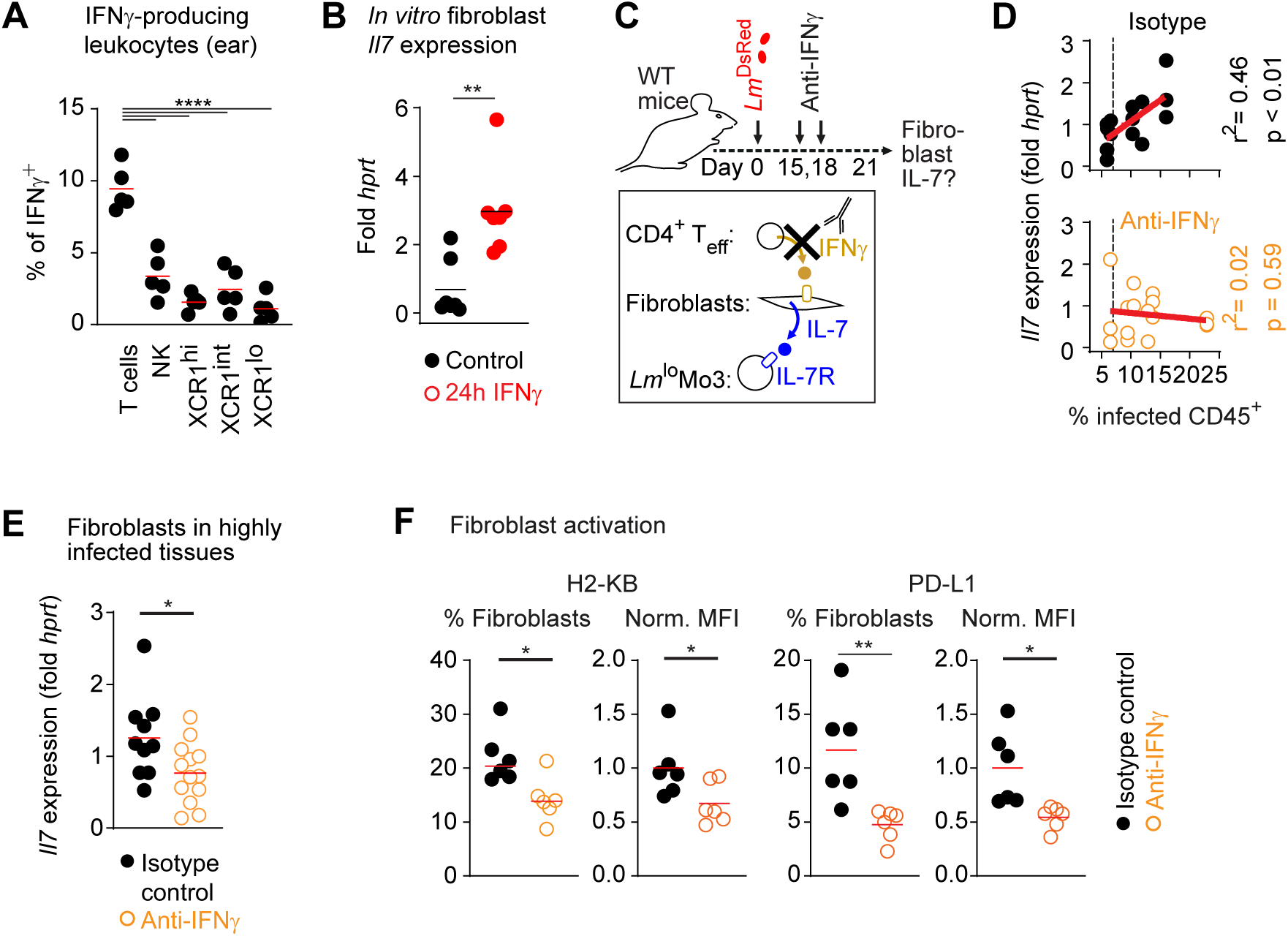
IFNγ activates fibroblasts to produce IL-7 depending of the infection amplitude. **A**. Proportion of IFNγ^+^ produced by T, NK, and dendritic cells at week 3 p.i. Each symbol represents one individual mouse ear. Horizontal lines denote the mean. ****, p<0.0001 according to unpaired t-test. **B.** Expression of the *Il7* cytokine gene in ex vivo cultivated fibroblasts without (black symbols) and with (red symbols) addition of IFNγ^+^ for 24h. Horizontal lines denote the median. **, p<0.01 according to Mann-Whitney test. **C.** Experimental setup to test the impact of anti-IFNγ treatment in fibroblasts within an ongoing *L. major* infection. **D.** Correlation (r^2^, with p value indicated) and linear regression (red line) between *L. major* infection rate (determined by flow cytometry) and *Il7* expression level (qPCR) in fibroblasts within the infected ear (week 3 p.i.), in either isotype control (black symbols, upper graph) or anti-IFNγ treated (orange symbols, lower graph) animals. Each symbol represents one replicate of flow cytometry analysis from six individual infected ears per condition with qPCR of up to 3 pools of 200 fibroblasts sorted from the same infected ear. The vertical dashed line shows the threshold defined for analyzing highly infected tissues. **E.** Comparison of *Il7* expression in fibroblasts sorted from tissues with an infection rate above the threshold (dashed line) shown in (D). **F.** H2KB^+^ or PD-L1^+^ fraction and H2KB or PD-L1 MFI among fibroblasts isolated from week 3 p.i. infected ears of animals treated with either isotype control (black symbols) or anti-IFNγ antibody (orange symbols). Each symbol in (E-G) represents one individual mouse ear, horizontal lines denote the mean. ***, p<0.001; **, p<0.01; *, p<0.05 according to unpaired t-test. Data were collected in two independent experiments.

In order to demonstrate the impact of IFNγ signaling on IL-7 production by fibroblasts in vivo, we treated *L. major*-infected mice with IFNγ-specific blocking or isotype-matched control antibodies (Fig. 5C). In fibroblasts from the infected tissue of control mice, we observed a strong correlation between *Il7* gene expression and the cellular infection rate. In contrast, this correlation was lost after anti-IFNγ treatment (Fig. 5D). Consequently, anti-IFNγ treatment reduced fibroblasts *Il7* expression in strongly infected tissues (Fig. 5E), and decreased fibroblast activation (Fig. 5F and S6C). Therefore, we concluded that the IFNγ produced during the T cell response against *L. major* induces IL-7 production in fibroblasts, thereby activating immunosuppressive PDPN^+^IL-7R^+^ *Lm*^lo^Mo3 which subsequently blunt pathogen clearance.

### Anti-IL-7/IL-7R treatment results in an enhanced immune response against *L. major*

Our data suggested that targeting IL-7R signaling might offer a possibility to prevent immunosuppression by PDPN^+^IL-7R^+^ *Lm*^lo^Mo3, thus enhancing effector T cell responses and reducing pathogen burden. Sustained long-term treatment with anti-IL-7R antibody has been shown to impair T cell responses (Colpitts et al., 2009). However, IL-7R has been reported to be critical for maintaining mainly naïve and memory T cells, but less important for effector T cells (Capitini et al., 2009; Fry and Mackall, 2002; Huster et al., 2004). Thus, IL-7/IL-7R-blockade could suspend immunosuppression by PDPN^+^IL-7R^+^ *Lm*^lo^Mo3 when applied temporarily during the acute infection.

To test if such transient anti-IL-7/IL-7R treatment improves the control of *L. major* infections, we injected *L. major*^DsRed^-infected mice with an anti-IL-7/IL-7R antibody cocktail after the establishment of infection (Fig. 6A). As expected for IL-7-depleting conditions (Grabstein et al., 1993), the treatment resulted in a decline of immature B cells in the BM and spleen, but not in mature B cells (Fig. S7A-B) three days after antibody injection. We also observed a slight reduction in total CD4^+^ T cells, while the numbers of central memory, effector or regulatory T cells were unchanged in the draining lymph node (Fig. S7C). This indicated that the short-term treatment did not profoundly affect the peripheral lymphocyte homeostasis, or T cell effector populations. Importantly, anti-IL-7/IL-7R treatment resulted within 3-5 days in strongly reduced *L. major* infection (Fig. 6B-C), enhanced IFNγ production by effector T cells (Fig. 6D and S7D), and increased fibroblast activation (Fig.6E and S7E) in the skin. To explore whether anti-IL-7/IL-7R blockade also increased the induction of IFNγ-dependent antimicrobial effector functions (Muller et al., 2012), we determined iNOS expression in monocyte-derived cells isolated from the site of infection and found that anti-IL-7/IL-7R treatment resulted in higher iNOS expression in these cells (Fig.6F and S7F). Collectively, these data indicated that IL-7/IL-7R blockade efficiently increased the adaptive immune response against established skin infections with *L. major*. In order to address whether impaired IL-7R signaling had a long-term effect on pathogen control, and immune activation, we analysed the impact of anti-IL-7/IL-7R treatment several weeks after antibody treatment. For this, *L. major*^DsRed^ infected mice were injected with anti-IL-7/IL-7R antibody or isotype control at day 16 and 20 p.i., and analyzed at week 7 p.i. for pathogen control and immune activation (Fig. 6G). Weekly measurement of the infected ear thickness indicated a transiently higher inflammation in anti-IL-7/IL-7R injected versus control animals, suggesting that the treatment increased the amplitude of the immune response (Fig. 6F). At week 7 p.i., *L. major* tissue burden was reduced compared to isotype control-treated animals (Fig. 6I), indicating that although the local pathology seemed comparable at this late time point, the transient anti-IL-7/IL-7R had a lasting effect on pathogen control. Furthermore, the fraction of IFNγ-producing effector T cells was strongly increased in the draining lymph nodes of anti-IL-7/IL-7R treated animals (Fig. 6J and S7G). These data suggested that the transient blockade of IL-7/IL-7R and suspension of the immunosuppressive function of *Lm*^lo^Mo3 in the early phase of the response supported a long-lived CD4^+^ T cell response and pathogen control.

**Figure 6.**
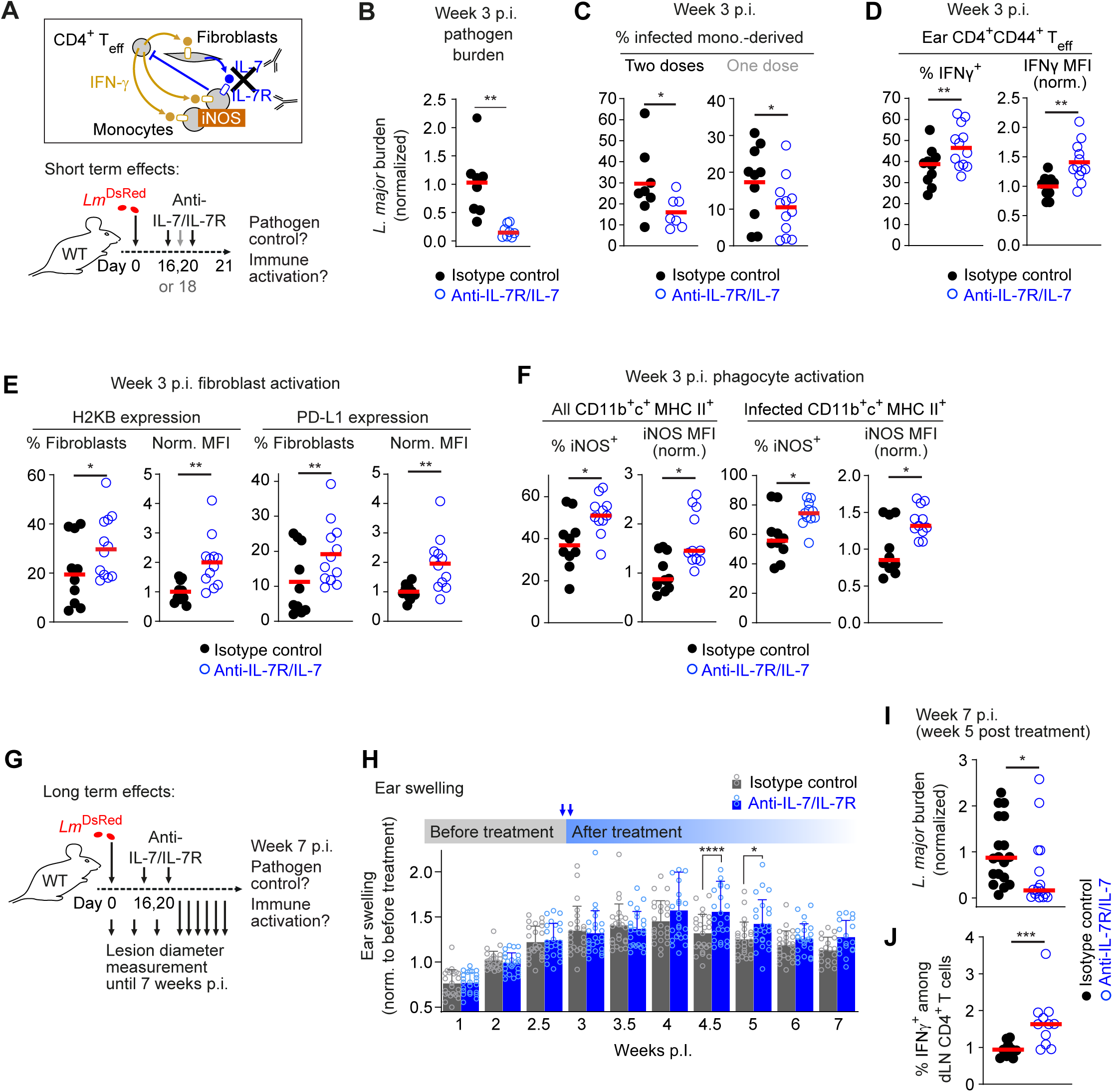
Anti-IL-7/IL-7R treatment results in an enhanced immune response against *L. major***. A.** Hypothesis (box) and experimental setup for testing the effects of anti-IL-7/IL-7R treatment during *L. major* infection. Treatment was applied twice (day 16, 20, black arrows, data shown in panel B-F) or once (day 18, grey arrow, data shown in panel C). **B-C.** Limiting dilution assay (LDA) of *L. major* tissue burden in the ear following anti-IL-7/IL-7R treatment normalized to isotype control treated animals (B) and *L. major* infected cell fraction among recruited monocytes with two-dosage (left panel) or single-dosage anti-IL-7/IL-7R treatment (right panel) (C). **D.** IFNγ^+^ fraction (left) and IFNγ MFI (normalized to isotype control treatment, right) among effector CD4^+^ T cells isolated from the ears at week 3 p.i. of either isotype antibody (black symbols) or anti-IL-7R/IL-7 treated animals (blue symbols). **E.** H2KB^+^ and PD-L1^+^ fraction among fibroblasts, and the MFI of H2KB or PD-L1 among fibroblasts (normalized to isotype control animals), either isotype control (black symbols) or anti-IL-7R/IL-7 treated (blue symbols). **F.** iNOS^+^ fraction and iNOS MFI (normalized to isotype control animals) among total recruited monocyte-derived cells (left panel) and infected monocyte-derived cells (right panel), either isotype control (black symbols) or anti-IL-7R/IL-7 treated samples (blue symbols). Each symbol in (B-F) represents one individual mouse ear, data pooled from two independent experiments. Horizontal lines denote the mean. **, p<0.01; *, p<0.05 according to unpaired t-test. **G**. Experimental setup to determine the long-term effects of anti-IL-7/IL-7R treatment. **H.** Infection-associated immune cell recruitment determined by ear diameter of isotype control-treated (grey) and anti-IL-7/IL-7R treated (blue) animals. Data were normalized to the mean of the corresponding values of the same ear measured before the treatment. Bars denote the mean+SD. Each symbol represents one individual ear repeatedly measured throughout the infection time course. Data obtained from two independent experiments. ****, p<0.0001 *, p<0.05 according to 2-way ANOVA (treatment, experiment) with Bonferroni post-test. **I.** Limiting dilution assay (LDA) of *L. major* tissue burden in the ear at week 7 p.i. of animals previously treated (week 3 p.i.) with isotype control-treated (black symbols) or anti-IL-7/IL-7R treated (blue symbols). **J.** IFNγ^+^ fraction among CD4^+^ T cells isolated from the draining lymph node at week 7 p.i. of animals previously treated (week 3 p.i.) with either isotype control (black symbols) or anti-IL-7R/IL-7 antibody (blue symbols). Each symbol in (I-J) represents one individual ear. Data pooled from two independent experiments and normalized to control animals of each experiment for limiting dilution. Horizontal lines denote the median. *, p<0.05 according unpaired Mann-Whitney test. Each symbol represents one individual mouse ear. Horizontal lines denote the mean. ***, p<0.001; according to unpaired Mann-Whitney test.

Altogether, our data therefore indicate that the IL-7 produced by fibroblasts in response to IFNγ drives PDPN⁺IL-7R⁺ *Lm*^lo^Mo3 cells blunting CD4⁺ T-cell effector functions, and targeted manipulation of this pathway can facilitate robust anti-*Leishmania major* immunity.

## Discussion

Mechanisms that restrict overshooting effector functions are an important component of immune responses to prevent immunopathological tissue damage (Formaglio et al., 2021; Mishra et al., 2013; Postat et al., 2018; Sugimoto et al., 2016). As such, the effects of IFNγ, a main mediator of cellular immune responses, are controlled on different levels, including the restriction of IFNγ-producing cells (Badovinac et al., 2003) and inhibition of IFNγ signaling (Alexander et al., 1999; Marine et al., 1999). Pathogens can benefit from such regulatory mechanisms to establish efficient infection (Collins-McMillen et al., 2017; Dolgachev et al., 2014; Drewry et al., 2019; Mariotti et al., 2004; Navarro et al., 2011), and also immune responses against tumors can be hampered by mechanisms that restrict IFNγ function (Janson et al., 2008). A better understanding of the cellular and molecular interactions underlying these immunosuppressive functions would greatly facilitate the development of novel immunotherapeutic approaches (Gocher et al., 2022; Wykes and Lewin, 2018). Such approaches have so far mainly focused on T cell functions directly, especially for the treatment of cancer (Ellis et al., 2021), while the potential of monocyte-derived functions is only beginning to be acknowledged (Kruse et al., 2023; Mantovani et al., 2022). We describe here a new self-limiting mechanism by which increasing IFNγ levels initiate a feedback loop that ultimately prompts IL-7R expressing monocyte-derived cells to prevent effector T cells from producing more IFNγ. Interruption of this feedback loop increased effector T cell function and promoted pathogen clearance.

Our unbiased identification of a PDPN⁺IL-7R⁺ monocyte-derived subset that harbors low-proliferating *L. major* and suppresses CD4⁺ T-cell effector function adds a new layer to the picture of myeloid cell heterogeneity during chronic infection. The plasticity of myeloid cells at infection sites has been intensely studied, with ample evidence for immunomodulatory functions, for example PD-L2^+^CD206^+^ monocyte-derived macrophages in the context of helminth infection (Gundra et al., 2017; Gundra et al., 2014; Oyesola et al., 2023), or CD206^+^ tissue resident macrophages which promote type 2 immunity in non-healing *L. major* Seidman strain infection (Lee et al., 2018; Lee et al., 2020; Lee et al., 2023). Furthermore, CCR2^+^ monocytes have been shown adopt a dual role in the establishment of parasite-permissive conditions and the activation of parasite control depending on their time point of recruitment and exposure to the local cytokine milieu (Carneiro et al., 2020). Our data extend these concepts by demonstrating that, downstream of IFNγ, stromal fibroblasts secrete IL-7 that engages IL-7R on a PDPN⁺ monocyte-derived population, driving these cells towards a hypo-inflammatory phenotype which dampens CD4^+^ T cell responses.

We clearly show that PDPN^+^IL-7R^+^ *Lm*^lo^Mo3 cells originate from CCR2^+^ monocytes and have a turnover rate of far more than five days, and do not express TRM markers such as CD206. Thus, we conclude that the functional overlap with alternatively activated CD206^+^ tissue-resident macrophages described previously is at most partial, and that the PDPN^+^IL-7R^+^ *Lm*^lo^Mo3 adopt their phenotype upon exposure to the infection site. In line with this, previous reports indicate that both macrophages and monocytes can upregulate cell surface expression of PDPN and IL-7R in response to inflammatory stimuli (Hou et al., 2010; Kerrigan et al., 2012; Yazdani et al., 2024).

IL-7R expression by myeloid cells has been shown to promote the development of tissue-resident macrophages in early ontogeny in mice (Cool et al., 2020; Leung et al., 2019). In humans, monocyte-derived lung macrophages have been reported to express IL-7R during their development under non-inflammatory conditions (Evren et al., 2021), and upregulate IL-7R expression in response to inflammation (Al-Mossawi et al., 2019; Hu et al., 2008). Consequently, IL-7R expressing myeloid cells were observed in a variety of inflammatory and infectious diseases. For example, serum from patients infected with *Mycobacterium tuberculosis* (Mtb) as well as Mtb products induces IL-7R on monocytes (Harelimana et al., 2022). Furthermore, a role of IL-7R in monocytes in modulating the immune response has been suggested in human patients of SARS COV2 infection, where their presence in the blood was correlated with lower disease pathology (Zhang et al., 2022). Our analysis of skin melanoma tissue and re-analysis of published tumor-associated scRNAseq data suggests the presence of PDPN^+^IL-7R^+^ hypo-inflammatory monocyte-derived cells also in the inflamed tumor microenvironment. In line with this, *PDPN* and *IL7R* expression was found in monocyte-derived subpopulations of human head and neck carcinoma (Hoffmann et al., 2022), and *IL7R* was found to be a signature gene for tumor-associated monocyte-derived cell subsets in a variety of malignancies (Cheng et al., 2021; Mulder et al., 2021; Nalio Ramos et al., 2022). Altogether, these findings suggest a more general role for IL-7R expressing monocyte-derived cells in dampening immune responses in a variety of diseases. With the identification of a feedback loop fueled by IFNγ, which induces IL-7 production by activated fibroblasts, we provide a multicellular interaction network for the immunosuppressive function of PDPN^+^IL-7R^+^ *Lm*^lo^Mo3. Of note, we also observed PDPN^+^IL-7R^-^ single-positive monocyte-derived cells. These cells might represent an intermediate state, but, alternatively, might deploy immunoregulatory functions independently of the described IL-7 dependent feedback switch, since not only IL-7R, but also PDPN has been shown to be involved in immunoregulatory mechanisms (Xie et al., 2020).

The enhancement of pathogen clearance in response to i) *Il7r* deficiency in or depletion of PDPN^+^IL-7R^+^ *Lm*^lo^Mo3, ii) fibroblast-specific *Il7* gene inactivation, and iii) IL-7/IL-7R blockade, strongly suggests that IL-7 can mediate an immunosuppressive function. While other cell types could also produce IFNγ to enhance pathogen control, T cells seem to be a main and sufficient IFNγ producer for iNOS induction (Muller et al., 2012), and thereby a very likely target of the described feedback mechanism. At first sight this appears counter-intuitive, since IL-7 stimulates T cell immunity at multiple levels, such as thymic T cell development and the survival of naïve and memory T cells in the periphery. However, this is not the case for effector T cells, which down-regulate IL-7R expression and are thus insensitive to direct IL-7 effects (Bachmann et al., 2005; Huster et al., 2004; Kaech et al., 2003). Importantly, endogenous IL-7 had previously been shown to suppress IFNγ production and to exacerbate disease in a skin infection model of mice with *Schistosoma mansoni* (Wolowczuk et al., 1997). Moreover, treatment with recombinant IL-7 can aggravate *L. major* infection in susceptible mice (Gessner et al., 1995). In line with these reports, we found that short-term blockade of IL-7/IL-7R affects the number of naïve T and B cells in secondary lymphoid organs, but not of effector T cells, whose IFNγ production is increased, resulting in enhanced control of the pathogen. We have shown previously that even slight manipulation of pathogen permissiveness at week 3 p.i. results in enhanced control later on (Formaglio et al., 2021). In line with this, we observed an altered course of pathology, with stronger ear swelling 2-3 weeks after anti-IL-7/IL-7R treatment. The treatment also resulted in more efficient pathogen clearance on the long-term perspective, including stronger IFNγ production by T cells, which indicates a long-lasting effect of the transient anti-IL-7/IL-7R administration at the peak of infection. To specify the critical source of IL-7 mediating an immunosuppressing effect, we employed a conditional *Il7* knockout mediated by *Prrx-cre*, which is active in the majority of skin fibroblasts around birth (Leavitt et al., 2020; Liu et al., 2022), and is upregulated during wound healing (Lee et al., 2022). Importantly, we show that *Il7* expression is affected by the conditional knockout only in fibroblasts, but not any other stromal cell population sorted from the skin. The finding that *Prrx-cre* mediated *Il7* was sufficient to enhance *L. major* clearance and effector CD4^+^ T cell function therefore suggests that fibroblasts are critical producers of IL-7 in the mechanism. Skin fibroblasts have been shown to increase *Il7* expression in response to inflammation in a variety of models, including the infection of the ear dermis with *L. major* (Peduto et al., 2009), which we could confirm. IFNγ, which we could show to be mainly produced by T cells in our system, is sufficient to induce *Il7* expression upon injection in the skin of uninfected mice. Conversely, anti-IFNγ injection decreases *Il7* expression in a model of autoimmune alopecia (Dai et al., 2021). Therefore, it is conceivable that IFNγ produced by *L. major*-specific T cells (Muller et al., 2012) initiates an IL-7 mediated feedback from fibroblasts which ultimately dampens pathogen clearance. This mechanism might constitute a feedback switch that activates immunosuppressive monocyte-derived cells as soon as local IFNγ production in the course of the T cell response exceeds a certain threshold.

Targeting immunoregulatory mechanisms carries the risk of inducing immune-related adverse events (Yan et al., 2024), including pathological effects of unrestricted T cell functions (Byrne and Fisher, 2017; Johnson et al., 2016; Murakami et al., 2016). Of note, we did only observe a slight, transient increase of inflammation, while pathogen control was long-lasting. Two explanations are likely for this finding: First, efficient pathogen control could decrease pathogen burden-dependent inflammation (Formaglio et al., 2021), thereby masking the increased immune activation related to impaired IL-7 signaling. Second, the effector molecule nitric oxide, which is likely produced at higher amounts in response to the increased IFNγ, can restrict leukocyte recruitment (Mishra et al., 2013; Postat et al., 2018), thus counteracting an even more severe pathology in the anti-IL-7/IL-7R treated animals. Therefore, targeting IL-7 mediated immunosuppression might be especially well-suited for enhancing immune reactions involving nitric oxide-mediated killing by monocyte-derived cells, as nitric oxide seems to provide an additional layer of immunoregulation. Such monocyte-mediated killing has been demonstrated not only in immunity against intracellular pathogens (Olekhnovitch et al., 2014), but also during T cell therapy against cancer (Kruse et al., 2023).

Taken together, we describe a cytokine-dependent feedback loop that can restrict T cell activation in response to excessive IFNγ production, a mechanism from which the pathogen can benefit to perpetuate the infection. Targeting this mechanism to adjust the activation of effector T cell functions opens new avenues for future immunotherapeutic approaches.

## Methods

### Ethics Statement

All animal experiments were reviewed and approved by the Ethics Committee of the Office for Veterinary Affairs of the State of Saxony-Anhalt, Germany (permit license numbers 42502-2-1314 UNI MD, 42502-2-1393 Uni MD, 42502-2-1575 Uni MD, 42502-2-1586 Uni MD, 42502-2-1615 Uni MD, 42502-2-1672 Uni MD, and 42502-2-1735 Uni MD) in accordance with the legislation of both the European Union (Council Directive 499 2010/63/EU) and the Federal Republic of Germany (according to § 8, Section 1 TierSchG, and TierSchVersV).

#### Parasites, tumor and mouse infections

*L. major* LRC-L137 V121 wild-type, DsRed-, and mKikume-expressing (*Lm*^SWITCH^) parasites were described previously (Muller et al., 2013; Sorensen et al., 2003). *L. major* expressing the cyan fluorescent protein mTurquoise (*Lm*^Turquoise^) was generated by targeted integration of the mTurquoise gene into the recombinant RNA locus of L. major LRC-L137 V121. For this, the mTurquoise gene (Goedhart et al., 2010) sequence was synthesized *de novo* (Eurofins Scientific) and cloned via BamHI-BglII and NotI (New England Biolabs) into a pLEXSY-Hyg2 vector (Jena Bioscience). The resulting plasmid was linearized usin SwaI (New England Biolabs) and electroporated into *L. major*. Stable transfectants were selected with 30 mg/ml hygromycin B (Invitrogen). Single clones were obtained by limiting dilution. All parasites were grown in M119 medium completed with 10% heat-inactivated fetal calf serum, 0.1 mM adenine, 1 mg/ml biotin, 5 mg/ml hemin, and 2 mg/ml biopterin (all from Sigma) and kept for a maximum of six passages.

Wild-type C57BL/6J and CD45.1 (B6.SJL-*Ptprc^a^Pepc^b^*/BoyJ, JAX stock #002014) mice, *Il7r^-/-^*(B6.129S7-*Il7r^tm1Imx^*/J, JAX stock #002295,) (Peschon et al., 1994), *Rag1^-/-^* (B6.129S7-*Rag1^tm1Mom^*/J, JAX stock #002096) (Mombaerts et al., 1992), and mKikumeGR (Tg(CAG-KikGR)33Hadj/J, JAX stock #013753) (Nowotschin and Hadjantonakis, 2009) mice were purchased from The Jackson Laboratory (Bar Harbor, ME) via Charles River (Sulzfeld, Germany). Bone marrow from Ccr2-creERT2 (C57BL/6NTac-*Ccr2^tm2982(T2A-Cre7ESR1-T2A-mKate2)BB^*) (Croxford et al., 2015) x Ai14(B6.Cg-*Gt(ROSA)26Sor^tm14(CAG-tdTomato)Hze^*) (Madisen et al., 2010) mice was provided by Melanie Greter and Burkhard Becher (University of Zürich, Switzerland). To obtain *Prrx-cre* × *Il7^fl/fl^*animals, *Prrx-cre* (B6.Cg-Tg(Prrx1-cre)1Cjt/J, JAX stock #005584) (Logan et al., 2002) mice were purchased from The Jackson Laboratory and crossed to mice harboring floxed *Il7* alleles (*Il7^fl^Il7^fl^*), which had been previously generated by removing a FRT flanked part of a “knockout-first” construct *Il7^tm1a(EUCOMM)Wtsi^* (provided by The European Conditional Mouse Mutagenesis Program (EUCOMM)) using a Flpo-transgenic (Kranz et al., 2010) cross (Knop et al., 2020). To obtain *Pdpn-cre* × *Il7r ^fl/fl^* and *Pdpn-cre* × iDTR animals, the sperm of Pdpn-cre mice (C57BL/6N-Tg(Pdpn-icre)14Biat/Biat, EM#11673) (Onder et al., 2011) were purchased from European Mouse Mutant Archive (EMMA) and used in artificial insemination procedures with *Il7r ^fl/fl^*(Il7rtm1.1Asin/J, JAX stock #022143) (McCaughtry et al., 2012) females and ROSA26iDTR (CBy.B6-Gt(ROSA)26Sortm1(HBEGF)Awai/J) (Buch et al., 2005) females, respectively. The animals were housed and bred under specific-pathogen-free conditions in the central animal facility (ZTL) of the Medical Faculty at Otto-von-Guericke-Magdeburg.

For infection, 2×10^6^ stationary phase promastigotes were washed and resuspended in PBS and subsequently injected in 10 μl into the ear dermis. Where indicated, Tamoxifen (Sigma-Aldrich) was dissolved in olive oil (Sigma-Aldrich) at a concentration of 20 mg/ml and administered by i.p. injection (100 mg/kg) for five consecutive days, either five days before the infection, or between days 11 through 15 p.i. The mouse melanoma cell line HCmel12 had been originally established from a primary melanoma in the Hgf-Cd4k^R24C^ mouse model by serial transplantation as described previously (Bald et al., 2014; Kruse et al., 2023), and were cultured in complete Roswell Park Memorial Institute (RPMI) medium consisting of RPMI 1640 medium (Life Technologies) supplemented with 10% foetal calf serum (Biochrome), 2 mM L-glutamine, 10 mM non-essential amino acids, 1 mM HEPES (all from Life Technologies), 20 µM 2-mercoptoethanol (Sigma), 100 IU ml−1 penicillin and 100 µg ml−1 streptomycin (Invitrogen) in a humidified incubator with 5% CO_2_. For tumor inoculation, a total of 2×10^5^ cells were injected intracutaneously (i.c.) into the shaved flanks of mice with a 30G (0.3×13 mm) injection needle (BD). Tumor development was monitored by inspection and palpation. Mice were euthanized at 21 days post-tumor injection or before if tumors exceeded 15 mm mean diameter or when mice showed signs of sickness in adherence with the local ethical regulations.

#### Antibody treatment

For anti-IL-7/IL-7R application, mice were intraperitoneally (i.p.) injected with 500 µg of anti-mouse IL-7Rα (clone A7R34, Bio X Cell), which is non-depleting (Mai et al., 2014; Seddon et al., 2003), and 500 µg of anti-mouse IL7 (clone M25, Bio X Cell) per treatment (Table S6). Control mice were i.p. injected with 500 µg of rat IgG2a isotype control (clone 2A3, Bio X Cell) and 500 µg of mouse IgG2b isotype control (clone MPC-11, Bio X Cell) per treatment. For anti-IFNγ application, mice were i.p. injected with 500 µg of anti-mouse IFNγ (clone R4-6A2, Bio X Cell) per treatment, while control mice received 500 µg of rat IgG1 isotype control (clone HRPN, Bio X Cell) per treatment.

#### Adoptive T cell and BM transfer, irradiation and BM chimera generation

For adoptive transfer, CD4^+^ T cells were isolated from axillary, brachial, cervical, and inguinal lymph nodes as well as from the spleen using the CD4 isolation kit and negative selection MACS purification (Miltenyi Biotech, Bergisch Gladbach, Germany) according to the manufacturer’s instructions. 1 × 10^7^ T cells per recipient were injected intravenously. For BM isolation, BM cells were flushed from tibia and femur with ice-cold non-supplemented RPMI 1640 (PAN Biotec) and filtered through 100 micron cell strainers. Cells were washed with non-supplemented PBS, and 4 x 10^7^ cells (for BM reconstitution) per recipient were resuspended in PBS and injected intravenously into the tail vein. For generating BM chimeras or fully BM reconstituted mice, recipients were γ-irradiated with 9.5 gray before reconstitution and infected 8 weeks later.

#### Photoconversion for proliferation measurement and optical time-stamping

Photoconversion of *Lm*^SWITCH^ parasites and cells in the mouse ear was performed with a 405 nm wavelength, 665 mW/cm2 collimated high-power LED (Thorlabs). The ears of anesthetized mice were fixed and illuminated from each side for 30 seconds at a distance of 20 cm and were analyzed after 48h by flow cytometry.

#### Flow cytometry

Cells were harvested from digested ears as previously described (Heyde et al., 2018). Draining lymph nodes or spleen were dissociated on a 70 μm cell strainer, and cells were washed with PBS. Surface staining of cells was done by using the antibodies specified in table S6. For intracellular staining of IL-1 beta (Pro-form), IL-6, IL-12 p35, CXCL9, Eomes, Foxp3, GATA3, RORgt, T-bet and IFNγ, the cell suspension was fixed for 1hr at 4°C with fixation buffer (Biolegend), following permeabilization with Perm/Wash solution (Biolegend) according to the manufacturer’s instructions and stained with the antibodies listed in table S6 prior to staining and analyzed on a Fortessa flow cytometer (BD Biosciences). Samples with a content of less than 40% of CD11c-expressing cells among CD45^+^ were excluded from quantitative analysis. The mKikume fluorescence with/without photoconversion was read out at 561 nm excitation and 610/20 nm emission, or 488 nm excitation and 530/30 nm emission, respectively. An autofluorescence signal was recorded at 488 excitation and 695/40 nm emission. Data were analyzed using the FlowJo software (FlowJo, LLC). qPCR of sorted cells was performed as described in the supplemental materials and methods.

#### Single-cell index sorting and SmartSeq2 sequencing library preparation

For scRNAseq, mouse CD45 MicroBeads (Miltenyi Biotec) were used to isolate leukocytes prior to cell surface staining according to manufacturer’s instructions. In order to review the complete phenotype of every single cell and associate every phenotype with each sorted single cell, index sorting was performed on FACS ARIA III sorter (BD Biosciences). Single cells were sorted into different wells from a 96-well plate (Costar) filled with 4.65 µl of Buffer RLT Plus (Qiagen), 0.25 µl of SUPERase-In RNase-Inhibitor (Invitrogen), and 0.1 µl of ERCC spike-Mix1 (Invitrogen) pre-diluted 1:500000 in RNAse-free water. For cell-population RNAseq, 50-100 cells per subpopulation per sample were sorted into the wells. Sorted cells were immediately frozen and stored at-80 °C. Single-cell libraries were prepared using Smart-seq2 protocol as previously described (Picelli et al., 2014) with minor modifications: Following cDNA amplification, ACTB expression was measured in each cDNA library by qPCR. All cDNA libraries were diluted aiming to an equivalent ct value for ACTB (average = 21.11, SD = 1.05) and were subsequently used as input for a tagmentation-based protocol (Nextera XT, Illumina) using one quarter of the recommended volumes, 10 min for tagmentation at 55°C (Saliba et al., 2016). Libraries were quantified using Qubit dsDNA Hs Assay (Life Technologies) and fragment size of libraries was analyzed using QIAxcel DNA high resolution Kit (Qiagen) and Bioanalyzer High Sensitivity DNA kit (Agilent). Libraries were normalized to 2 nM and pooled. Particularly, 218 of single-cell libraries or 87 of cell-population libraries were sequenced for HiSeq PE200 (Illumina). Sequencing data was analysed as described in the supplemental materials and methods.

#### Processing of RNA sequencing data

Raw fastq files were processed with Cutadapt (v3.7) (Martin, 2011) to remove poly-A tails and adapter sequences from the SMART-seq and Nextera protocols. Low-quality bases (Q-score < 20) and reads left with less than 25 bases after trimming were removed. Sequencing data were mapped to a reference genome using STAR with default parameters (Dobin et al., 2013). The reference genome was built by combining the mouse (GRCm38) and *Leishmania major* (ASM272v2) genomes as well as spike-ins (ERCC92) and the Kikume Green-Red transcript sequences. Raw count matrices were generated from the mapping results after read count quantification with featureCounts (Subread, v1.6.3) (Liao et al., 2014). Two count matrices for 218 single cells and 87 cell-population libraries were created.

#### Quality control and normalization of scRNAseq data

Low-quality cells with more than 2% mitochondrial read counts, more than 20% ERCC spike-ins read counts or less than 1000 detected genes were discarded. Only mouse genes with at least 5 read counts detected in at least 10 cells were retained. Read counts for the 194 retained single-cell transcriptomes were normalized with scNorm (Bacher et al., 2017). Information about the plate for technical biases (*Condition*) and gene length (*WithinSample*) was considered during the normalization procedure.

#### Dimensional reduction and clustering of scRNAseq data

Downstream analysis of the single-cell dataset has been performed with the Seurat R package (v4.0) (Hao et al., 2021). The normalized read count matrix was natural log-transformed and the top 1000 highly variables genes were identified (*FindVariableFeatures*). For dimensional reduction, principal component analysis was performed (*RunPCA*) to capture the principal components (PCs) explaining the highest amount of variation in the dataset. Batch effects between mice were corrected with Harmony (v0.1.1). The top 8 PCs from the corrected PCA were projected into a 2D space by Uniform Manifold Approximation Projection (*RunUMAP*) for data visualization. In the uncorrected data, we found that cells from three infected ears were strongly separated from the rest of the cells by in UMAP 1 versus UMAP 2 (see Fig. S1E). MACS-purified cells from these infected ears revealed a substantially lower percentage of CD11c^+^ cells than the remaining samples in other ears. While this war corrected for scRNAseq, we excluded cells from samples with a CD11c^+^ cell content of less than 40% from experiments with functional readout.

A graph-based unsupervised machine learning procedure was used to identify groups of single cells with similar transcriptomes, i.e. clusters. The top 8 PCs were used to build a KNN graph (*FindNeighbors*; k = 10) followed by partition into clusters with the Leiden algorithm (Traag et al., 2019). We used the C implementation for R of the Leiden algorithm (leidenbase v0.1.3, Trapnell Lab) with a resolution set at 0.08. Clustering results were used to label cells on the UMAP.

#### Differential expression testing and pathway analysis for single-cell data

Gene markers of each cluster were identified by a Wilcoxon Rank-Sum Test with Seurat (*FindAllMarkers*). Only genes that were expressed in at least 10% of the cells from a given cluster, with an absolute log2 fold-change > 0.25 and a Bonferroni adjusted p-value < 0.05 were considered as differentially expressed.

Gene Set Enrichment Analysis (v4.3.2) was used for pathway analysis (Subramanian et al., 2005). DEG testing was repeated over all genes by a Wilcoxon Rank-Sum Test for each cluster versus all others through Seurat (*FindMarkers*). Gene-ranked lists were generated after converting all uncorrected p-values to-log10(p-value) * fold-change direction. Ranked lists were used as input for GSEA preranked with default parameters and Gene Ontology Biological Process gene sets. Pathways up-or down-regulated with FDR < 0.05 were considered significant.

#### Quantitative PCR analysis

For validation of DEG identified in the scRNAseq, 50 cells per sample were sorted from each subpopulation according to the gating strategy shown in fig. S1K and lysed in 10μl lysis buffer (RNeasy Plus lysis buffer, QIAGEN) containing 5% SUPERase-In RNase-Inhibitor 20U/μl (Invitrogen). RNA extraction and reverse transcription were performed using the Smart-seq2 protocol as previously described (Picelli et al., 2014). Primers used are specified in table S6.

For measuring *Il7* gene expression, total RNA was isolated from *L. major*-infected ear tissue using TRIzol reagent (Invitrogen) and then was reverse-transcribed using the High-Capacity cDNA Reverse Transcription Kit (Applied Biosystems) following the manufacturer’s instructions. In order to correct for the immune cell infiltrate which was expected to contain leukocytes expressing *Hprt*, but not *Il7*) *hprt* CT values were corrected according to the mean difference in RNA extracted.

For measuring the *Il7* and *Prrx1* gene expression in fibroblasts, LEC and BEC, three replicates of 200 cells each were sorted from each mouse ear. RNA extraction and reverse transcription was performed using the Smart-Seq2 protocol as described above. Quantitative PCR analysis was done using a SYBR Green Master Mix (Applied Biosystems).

To run the quantitative PCR, 40 cycles of 95 °C (15 s), and 60 °C (60 s) followed by a dissociation protocol 95 °C (15 s), 60 °C (20 s), and 95 °C (15 s) were performed on an ABI Prism 7000 Sequence detection system (Applied Biosystems). Technical triplicates were performed for each sample.

#### Limiting dilution assay

*L. major* burden within ear tissue was measured using a limiting dilution assay. Quadruplicate samples of each ear homogenate were serially diluted in 1:2 steps in *Leishmania* culture medium in 96-well plates and incubated at 26 °C for 14 days. The initial parasite burden was calculated from the median of the highest dilutions of the quadruplicates which exhibited parasite growth. In order to compensate experiment-to-experiment variation, the values were normalized to the control groups for pooled data.

#### Proliferation analysis

The proliferation index of *L. major* for flow cytometry was calculated based on the MFI of mKikumeGreen and mKikumeRed as described previously (Heyde et al., 2018). In brief, for visualizing qualitative comparisons within the same sample using the FlowJo software, values were plotted as

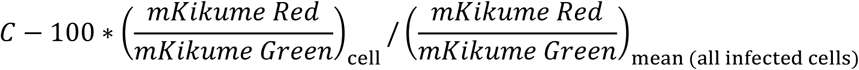

with C chosen between 100 and 250 and kept constant within the same sample for which the comparison was made, and the factor 100 was introduced to analyze integer fluorescence values in FlowJo.

For inter-sample comparison, the proliferation index was calculated as

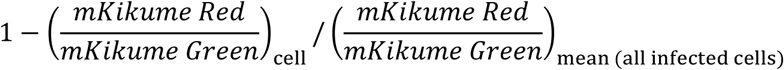

and represented as a percent deviation from the total infected cell population within one sample or imaged infection site.

#### Cell culture and stimulation

To generate a fibroblast culture, lymphoid stromal cells were isolated from peripheral lymph nodes of C57BL/6J mice as described previously (Knop et al., 2020). Afterwards, cells from four mice were pooled and seeded in a T25 flask in full medium (RPMI 1640 supplemented with 10% (v/v) FCS (FBS Superior; Sigma) /1% (v/v) P/S (Gibco)/2 mM L-glutamine (Gibco)/1 mM sodium pyruvate (Gibco)/0.1 mM HEPES (Gibco)/50 μM 2-mercaptoethanol (Sigma). Cells were cultured for 24h at 37°C and 5% CO_2_ and non-attached cells were removed. When reaching confluence, cells were detached using Trypsin/EDTA (Merck Biochrom) and seeded onto T75, and afterwards onto T175. Cells were detached using Trypsin/EDTA and treated with biotinylated anti-CD45 (30-F11, Biolegend) and anti-CD31 (MEC13.3, Biolegend)-specific antibodies in degassed staining buffer (PBS/ 2 mM EDTA (Carl Roth)/ 0,5% (w/v) BSA (AppliChem)) for 15min at 4°C. Cells were washed and resuspended in degassed PBS/ 2 mM EDTA and CD45+ and CD31+ cells were depleted using Streptavidin MicroBeads (Miltenyi Biotec), LS columns (Miltenyi Biotec) and MidiMACS separator (Miltenyi Biotec) according to manufacturer’s instructions. Cells were seeded in a T175 flask in full medium. When reaching confluence, cells were detached using Accutase (Stempro, Gibco) and stained with anti-CD45.2-APC (104, Biolegend), gp38-A488 (8.1.1, Biolegend) and CD31-PE/Cy7 (390, eBioscience)-specific antibodies for 30min at 4°C in staining buffer containing purified anti-CD16/32 (2.4G2 ATCC® HB-197TM). After washing with PBS/2 mM EDTA, cells were resuspended in full medium containing 2 μg/ml Ciprobay (Bayer) and CD45.2^-^ gp38^+^CD31^-^ cells were sorted using FACSARIA III (Becton Dickinson). The sorted cells were seeded at a density of 7400 cells/cm^2^ in full medium containing 2 μg/ml Ciprobay. After 48h, Ciprobay was removed and the fibroblasts were expanded until reaching confluence in a T175 flask.

For IFN-γ stimulation, cells were detached using Accutase and were seeded at 40000 cells/24-Well in 500 μl full medium without or with 50 ng/ml recombinant mouse IFN-γ (R&D Systems). After 24h, the cells were detached using Accutase and resuspended in TRIzol reagent (Invitrogen). RNA isolation, reverse transcription and RT-qPCR were performed as described previously (Knop et al., 2020). The following TaqMan® Gene Expression Assays (Thermo Fisher Scientific) were used according to the manufacturer’s instructions: Il7 (FAM-MGB probe Mm01295804 m1) and Hprt (FAM-MGB probe Mm00446968 m1). Each sample underwent analysis in triplicates on qTOWER3 G (Analytic Jena). Subsequently, relative quantification was performed according to the ΔCT method.

#### Re-analysis of published scRNAseq from tumor-associated monocyte-derived cells

Unsupervised Leiden clustering and differential gene expression analysis for tumor-associated monocyt-derived cells were obtained from our published data (Kruse et al., 2023) that had been deposited at the NCBI GEO under the accession GSE230427 (raw data) and GSE230427 (normalized and logarithmized count matrix). In brief, the data had been generated from individual tumors from which CD45^+^ cells were enriched using a positive selection kit (Miltenyi) and hashtagged with unique TotalSeq-B hashtag antibodies B0301-B0310 (1:300, Biolegend) and subsequently stained with fluorescently labeled antibodies. CD45^+^CD11b^+^Ly6G^-^ cells were sorted with an Aria III fluorescence-activated cell sorter (BD). Isolated cells were loaded onto one lane of a 10X Chromium microfluidics controller. Sequencing was performed on an Illumina Novaseq on two lanes of a S1 cartridge with 150 bp read length in paired end mode. Reading depth was calculated to obtain roughly 50,000 reads per cell for the gene expression library and 5,000 reads per cell for the hashtag library. Monocytes and macrophages from non-treated tumor clustered into T1 through T4 (Kruse et al., 2023) were re-analyzed regarding differential gene expression between cells classified as Cluster T4 versus Cluster T1 through T3, and cells classified as *Il7r*-expressing versus all other cells using a zero-inflated regression model from the R package MAST (Finak et al., 2015).

#### Statistical analysis

Statistical analysis was carried out with GraphPad Prism 8 (GraphPad Software, San Diego, CA, USA). To compare multiple samples in pairwise analysis within datasets with more than two experimental groups, one-way analysis of variance (ANOVA) was done for datasets that had passed a Shapiro-Wilk normal distribution test, Kruskal-Wallis tests were performed for datasets with non-normal distribution. Appropriate multiple comparison post-tests (Bonferroni’s for multiple comparisons, Dunnet’s for multiple pairwise comparisons or multiple comparisons with a control group in ANOVA, Dunn’s test for Kruskal-Wallis analyses) were employed as indicated in the respective figure legends. Two-group comparisons were analyzed using two-sided, unpaired or paired t-tests for data with normal distribution and Mann-Whitney tests for datasets for which a Shapiro-Wilk normal distribution suggested non-normal distribution. Representation of the median or mean, with standard deviation in cases where not all data points are shown individually, are indicated together with sample size in the figure legends.

## Acknowledgments

The helpful discussions with Tim Lämmermann, Tatjana Ruhl, Gabriel Sollberger, and all members of the Institute of Molecular and Clinical Immunology are gratefully acknowledged. We thank Tamas Dolowschiak for invaluable advice for setting up the scRNAseq library construction. We also thank Roland Hartig from the MPBIC platform and the team of the ZTL of the Medical Faculty OvGU for excellent technical support, and Ehsan Vafadarnejad for the support with initial scRNAseq data analysis.

## Author contributions

YF, TS, AES and AJM conceived of and designed the study. YF, IS, ER, LP, PG, LK, BK, LAD, VD, JA, NJN, MB, IB, JM, JP, CS, BP, and AJM conducted experiments, YF, ER, CT, LP, JM, SK, LK, BK, SF, AB, RG and AJM analyzed the data. NG, TT, BS, HVH, MG, BB, NJ, and TS provided critical technology. YF, TS, AES and AJM wrote the manuscript. All authors revised, commented and approved the manuscript.

## Data and materials availability

RNAseq data supporting the findings of this study have been deposited at Gene Expression Omnibus under accession no. GSE228455. R scripts used for the analysis of the scRNAseq data are available on GitHub: https://github.com/saliba-lab/Monocytes_leishmania

