## Supplemental figures S1-7 for "Fibroblast-IL-7 feedback drives PDPN^+^ monocytes to restrain CD4^+^ T cell responses"

**Supplemental Figure S1:** PDPN$^+$IL-7R$^+$ phagocytes infected with low proliferating *L. major* exhibit a hypo-inflammatory phenotype.

**Supplemental Figure S2:** Occurrence of PDPN$^+$IL-7R$^+$ monocyte-derived cells with a hypo-inflammatory expression profile in skin melanoma

**Supplemental Figure S3:** IL-7R$^+$PDPN$^+$ *Lm*$^{lo}$Mo3 develop from bone-marrow derived CCR2$^+$ monocytes and acquire their phenotype in the infected tissue.

**Supplemental Figure S4:** PDPN$^+$IL-7R$^+$ *Lm*$^{lo}$Mo3 dampen T cell responses against *L. major* infection *in vivo*.

**Supplemental Figure S5:** Fibroblast-produced IL-7 contributes to pathogen persistence and suppression of IFNγ production in effector T cells.

**Supplemental Figure S6:** IFNγ activates fibroblasts to produce IL-7 depending of the infection amplitude.

**Supplemental Figure S7:** Anti-IL-7/IL-7R treatment results in an enhanced immune response against *L. major*.

**Supplemental Figure S1: PDPN⁺IL-7R⁺ phagocytes infected with low proliferating *L. major* exhibit a hypo-inflammatory phenotype.**

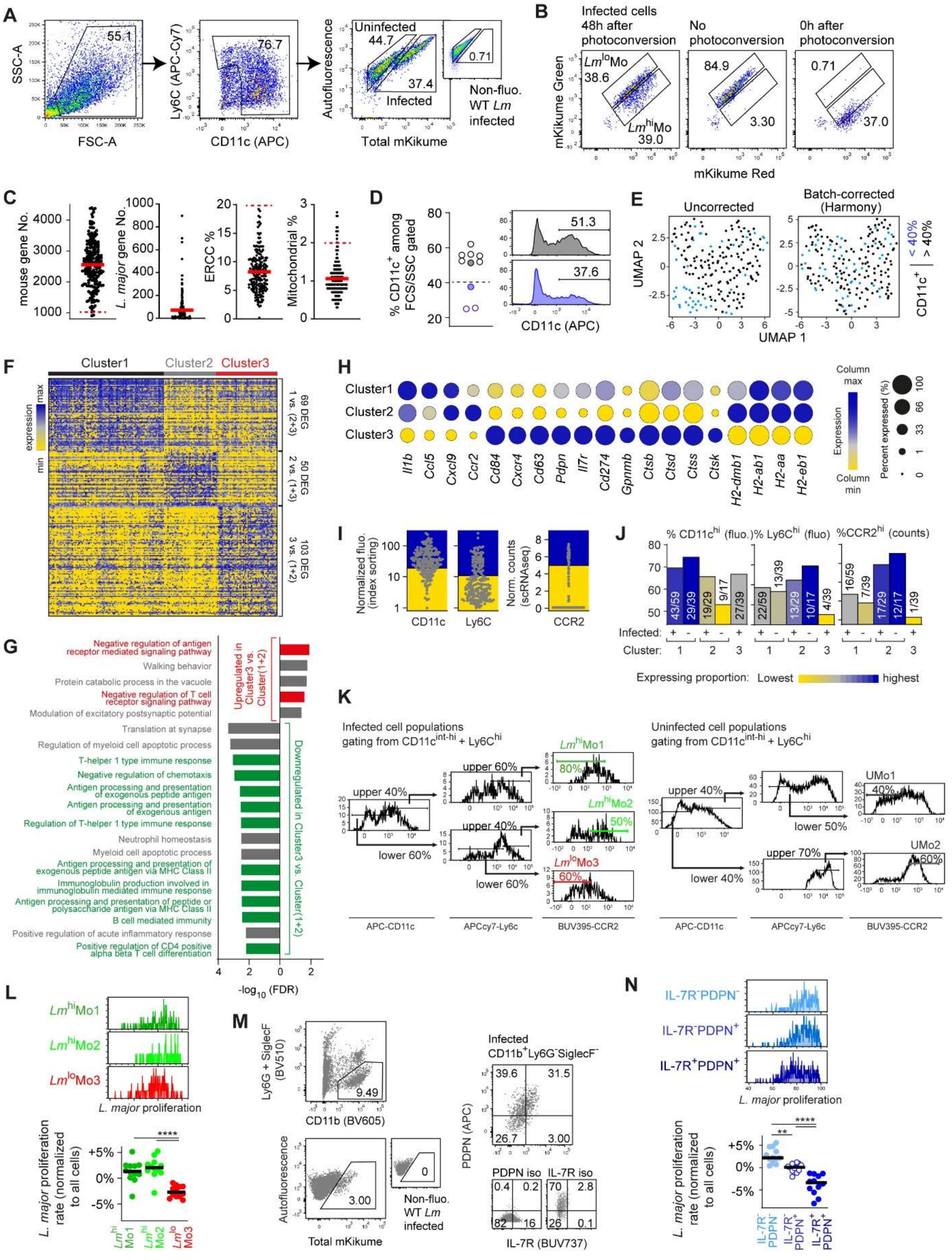


C57BL/6 mice were infected with $Lm^{SWITCH}$, parasites underwent photoconversion at 19 dpi and mice were analyzed at 21 dpi. **A.** Gating strategy for single-cell RNAseq of $Lm^{SWITCH}$-infected monocyte-derived cells. Non-fluorescent WT *L. major*-infected control is shown for comparison. **B.** Distinction of infected monocyte-derived cells according to the infection with high- and low-proliferating pathogens, based on their mKikume Green and Red fluorescence at 48h post photoconversion. Gating according to non-photoconverted (representing maximal proliferation) and photoconverted right before analysis (representing no proliferation). **C**. Basic statistics of scRNAseq including the number of mouse and *L. major* genes detected in each cell, and the fraction of ERCC control reads, or mitochondrial transcript reads among total reads of each sample. Horizontal lines denote the median. Dashed lines represent the thresholds for quality control. Data pooled from three independent experiments. **D.** $CD11c^+$ fraction among cells isolated by $CD45^+$-MACS of biological replicates of *L. major*-infected ears used for single-cell RNAseq. Each symbol represents one individual ear. Infected ears with a $CD11c^+$ fraction lower than 40% (blue) are indicated. The samples shown as histograms are shaded. Data pooled from three independent experiments. **E.** Uncorrected (left) and harmony batch-corrected (right) UMAP of transcriptomes of monocyte-derived cells sorted as shown in (A-B). Cells from infection sites with a $CD11c^+$ fraction lower than 40% are shown in blue. **F.** Heatmap of gene expression of DEG specifically upregulated in one of the three clusters with padj < 0.05. Details are in Extended Data Table S2. **G.** GO biological process term enrichment based on DEG found for Cluster 3. **H.** Normalized expression (color-code) and fraction of cells per population with detected expression (circle size) for selected DEG specifically expressed in cell cluster 1-3 identified in scRNAseq. **I.** Cutoff for high (blue) and low (yellow) expression of index markers CD11c and Ly6C (normalized as in (F)), and transcriptomic CCR2 expression level. Each dot represents one index-sorted single cell. **J.** Fraction of $CD11c^{hi}$, $Ly6C^{hi}$ and $CCR2^{hi}$ cells in each cluster, grouped according to infected and non-infected cells. **K.** Gating strategy for identification of $Lm^{hi}$Mo1, UMo1, $Lm^{hi}$Mo2, UMo2, and $Lm^{lo}$Mo3, independently of the proliferation reporter system based on CD11c, Ly6C and CCR2 expression and total *L. major* fluorescence. **L.** Examples (upper panel) and quantification of $Lm^{SWITCH}$ proliferation in $Lm^{hi}$Mo1, $Lm^{hi}$Mo2, and $Lm^{lo}$Mo3 gated according to the strategy shown in (K). Horizontal lines denote the median. Each dot represents one infected ear. ****, p<0.0001 according to Kruskal-Wallis multiple comparison with Dunn's post-test. **M.** Gating strategy for identification of $CD11b^+Ly6G^-SiglecF^-$ monocyte-derived cells according to IL-7R and PDPN expression. **N.** Examples (upper panel) and quantification of $Lm^{SWITCH}$

proliferation in $PDPN^+IL\text{-}7^+$, $PDPN^+IL\text{-}7^-$, and $PDPN^-IL\text{-}7^-$ monocyte-derived cells gated according to the strategy shown in (M). Horizontal lines denote the median. Each dot represents one infected ear. ****, $p<0.0001$; **, $p<0.01$ according to Kruskal-Wallis multiple comparison with Dunn's post-test.

**Supplemental Figure S2: Occurrence of IL-7R⁺PDPN⁺ monocyte-derived cells with a hypoinflammatory expression profile in skin melanoma**

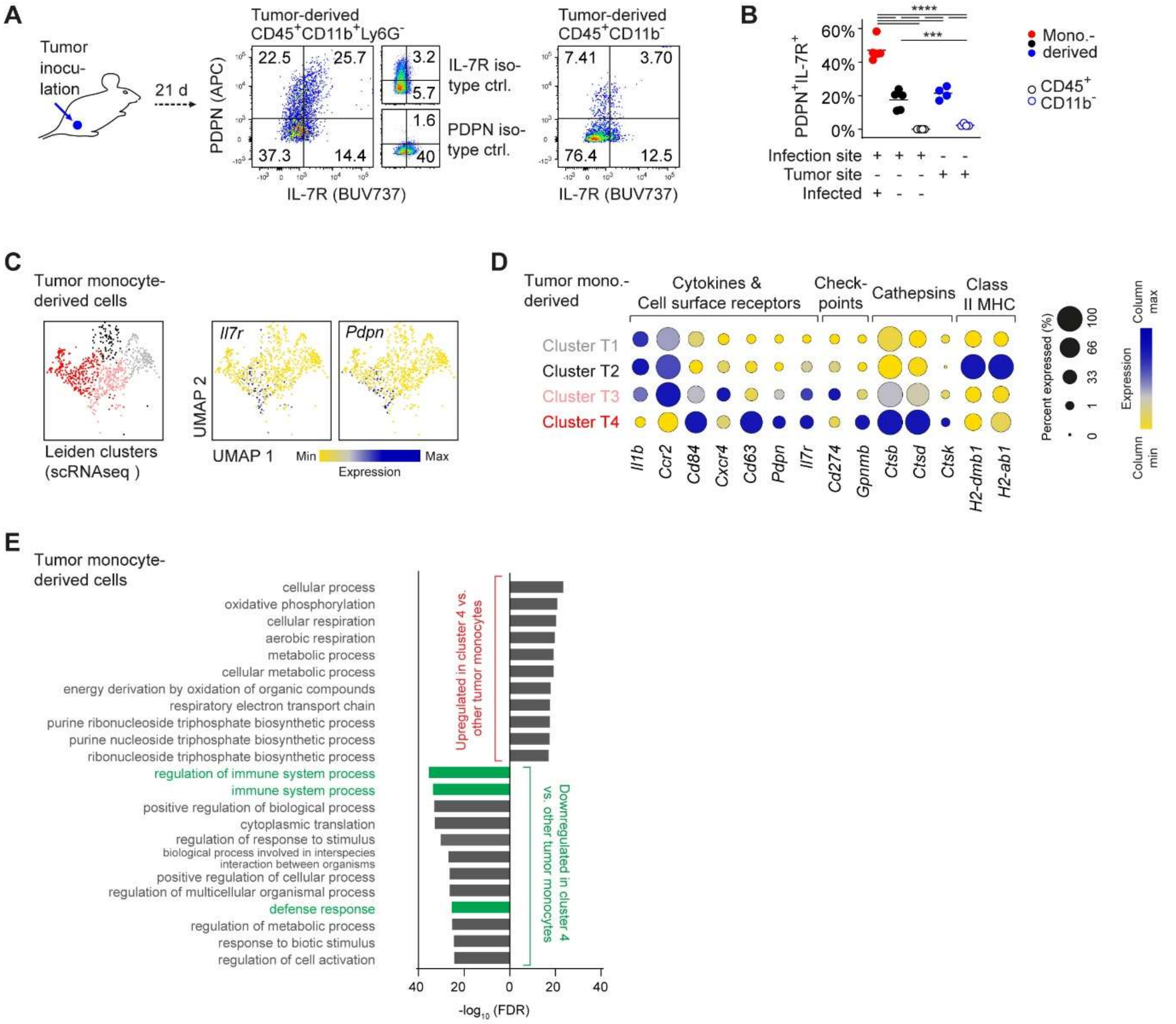


**A.** Experimental setup and gating strategy to analyze PDPN and IL-7R expression in CD45⁺CD11b⁺Ly6G⁻ monocyte-derived cells from Hcmel12 skin melanoma in C57BL/6 mice three weeks upon tumor inoculation. Applies to (B-E). Isotype controls and CD45⁺CD11b⁻ leukocytes are shown for comparison. **B.** Percentage of PDPN⁺IL-7R⁺ in CD45⁺CD11b⁺Ly6G⁻ monocyte-derived cells in infected (red dots) and uninfected (black dots) ear tissue or Hcmel12 tumors (blue dots) gated according to the gating strategy shown in A and figure S2F, respectively. CD45⁺CD11b⁻ leukocytes from the infection site (black) or tumor (blue) are shown as circles. Each dot represents one infected ear or one tumor analyzed. Horizontal lines denote the median. ****, $p<0.0001$; ***, $p<0.001$ according to Kruskal-Wallis multiple comparison with Dunn's post-test. **C.** UMAP representation, mapping and unsupervised Leiden clustering (left), normalized *Il7r* (middle) and *Pdpn* (right) expression of

CD45$^{+}$CD11b$^{+}$Ly6G$^{-}$ cells sorted from Hcmel12 skin melanoma deposited under NCBI GEO accession GSE230427 and GSE230427. **D.** Normalized expression (color-code) and fraction of cells in the tumor monocyte-derived cell cluster shown in (C) with detected expression (circle size) shown for DEG also found for *Lm*$^{lo}$Mo3 (see figure 1H). Only genes for which a significant difference between Cluster T4 versus the other clusters or between *Il7r* expressing versus *Il7r* non-expressing cells was found are shown (Table S4). **E.** GO biological process term enrichment based on DEG found for Cluster T4 versus all other clusters identified by unsupervised Leiden clustering of monocyte-derived cells in the tumor.

**Supplemental Figure S3: IL-7R⁺PDPN⁺ *Lm*^lo^Mo3 develop from bone-marrow derived CCR2⁺ monocytes and acquire their phenotype in the infected tissue.**

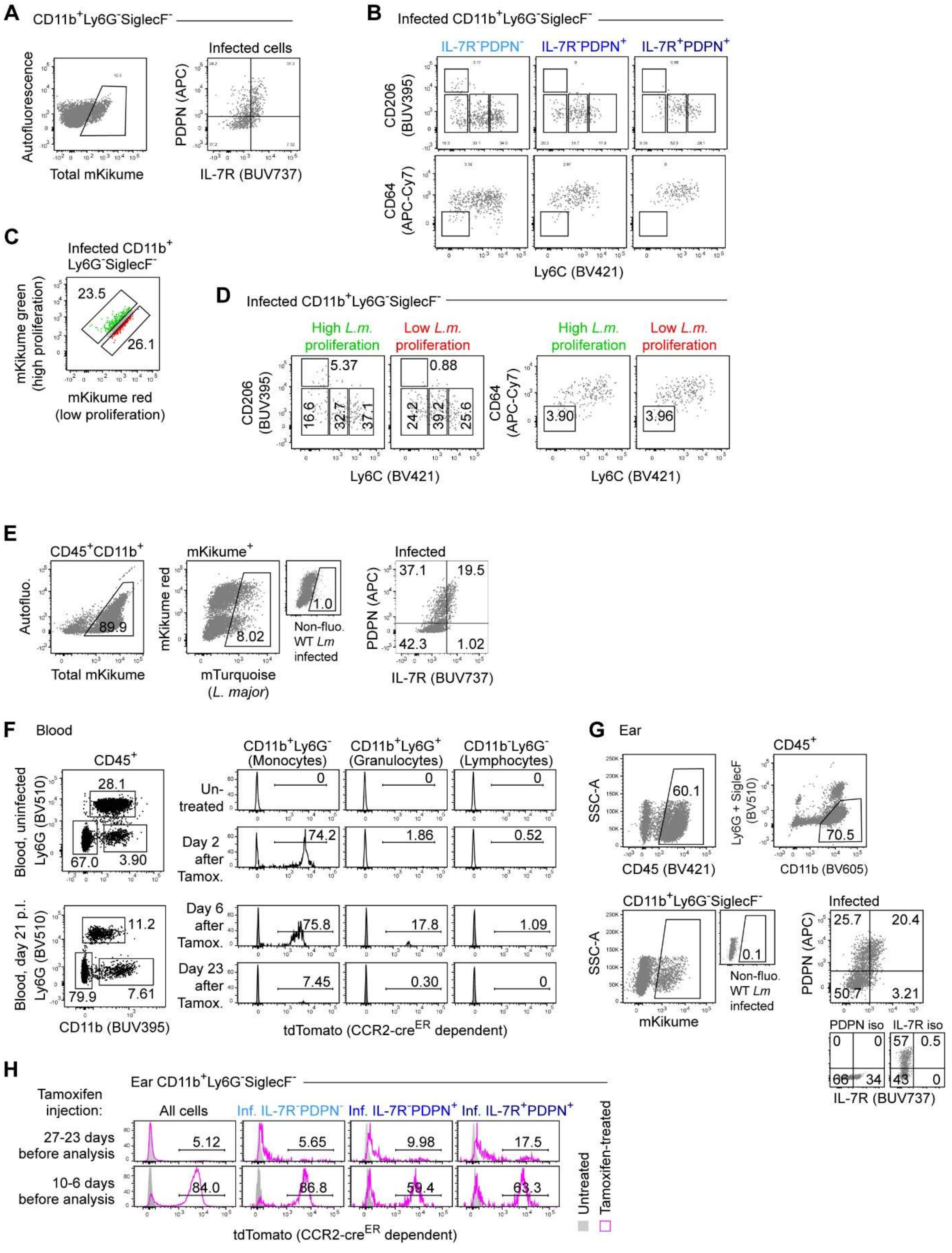


**A.** Gating on infected cells among CD11b⁺Ly6G⁻SiglecF⁻ monocyte-derived cells in *Lm*^SWITCH^-infected tissues. **B.** Distribution among tissue-resident and monocyte-derived dendritic cells (TR/MoDC), tissue-resident macrophages (TRM) and different monocyte-derived macrophage populations for CD11c$^{+}$Ly6G$^{-}$SiglecF$^{-}$ gated according to IL-7R and PDPN expression. **C.** Gating strategy to select CD11b$^{+}$Ly6G$^{-}$SiglecF$^{-}$ monocyte-derived cells infected with high- and low proliferating *Lm*$^{SWITCH}$. **D.** Distribution among tissue-resident and monocyte-derived dendritic cells (TR/MoDC), tissue-resident macrophages (TRM) and different monocyte-derived macrophage populations for CD11b$^{+}$Ly6G$^{-}$SiglecF$^{-}$ monocyte-derived cells infected with high- and low proliferating *Lm*$^{SWITCH}$. **E.** Gating on infected mKikume-expressing monocyte-derived cells infected with mTurquoise *L. major* according to IL-7R and PDPN expression. **F-H.** Ten weeks prior to *Lm*$^{SWITCH}$ infection, C57BL/6 mice were lethally irradiated and reconstituted with *CCR2-creER* × *LSL-tdTomato* BM (according to figure 2F). Mice were treated with Tamoxifen on the days indicated in figure 2J. **F.** TdTomato expression in blood CD45$^{+}$ cells before and day 2, 6 and 23 after the beginning of five daily Tamoxifen injections. **G.** Gating strategy to identify PDPN$^{-}$IL-7R$^{-}$, PDPN$^{+}$IL-7R$^{-}$ and PDPN$^{+}$IL-7R$^{+}$ monocyte-derived cell populations infected with mKikume-expressing *Lm*$^{SWITCH}$. **H.** Examples of tdTomato expression (magenta histograms) in infected PDPN$^{-}$IL-7R$^{-}$, PDPN$^{+}$IL-7R$^{-}$ and PDPN$^{+}$IL-7R$^{+}$ monocyte-derived cell populations at day 21 p.i., with Tamoxifen injection before infection (upper panels) or from day 16 p.i. for five days (lower panels). Grey histograms in (F) and (H) show non-transgenic control animals.

**Supplemental Figure S4: PDPN⁺IL-7R⁺ $Lm^{lo}$Mo3 dampen T cell responses against *L. major* infection *in vivo*.**

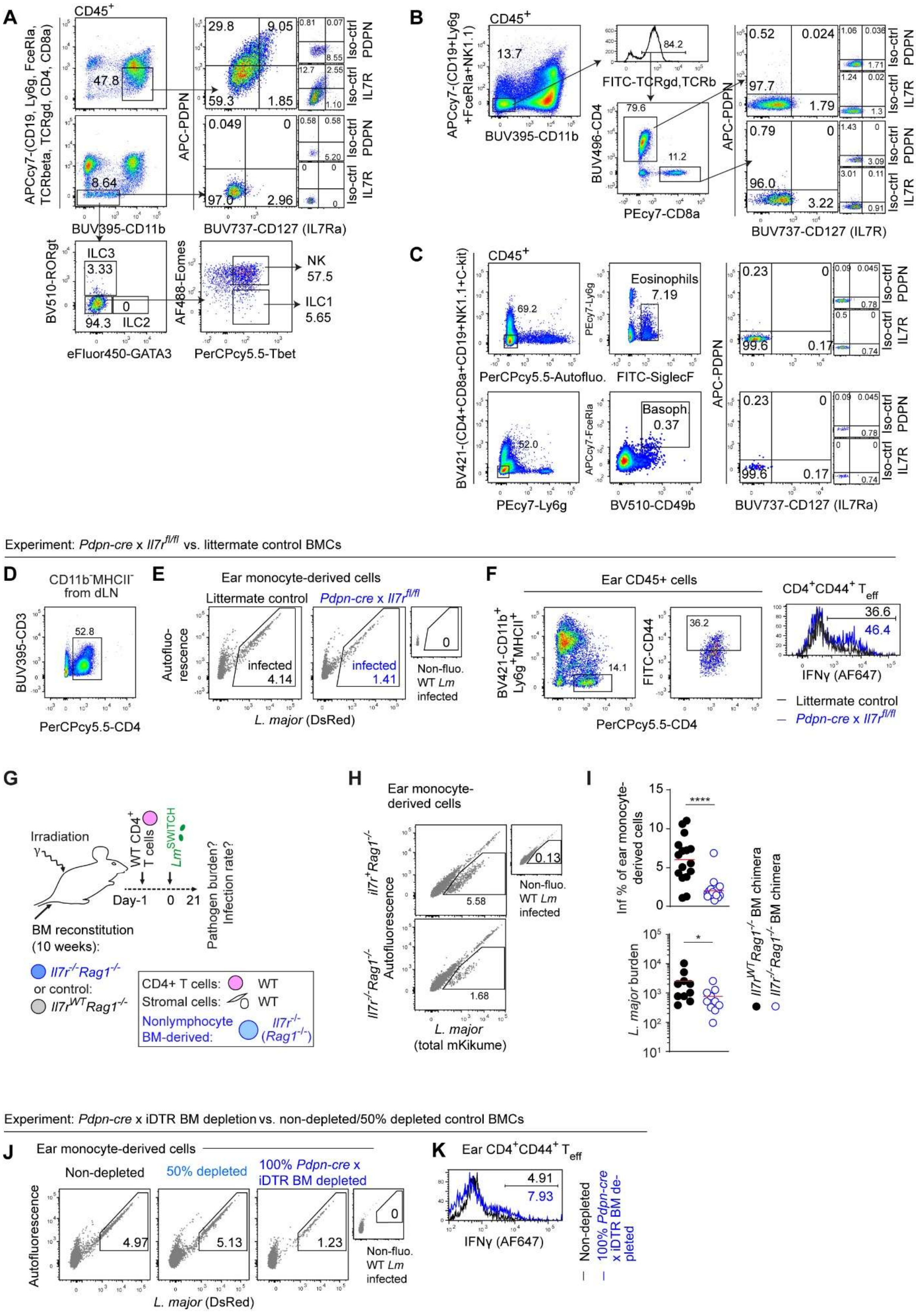

**A-C.** Gating strategies to determine PDPN and IL-7R expression in monocyte-derived cells, ILC1, ILC2, ILC3 and NK cells (A), $CD4^+$ and $CD8^+$ T cells (B), eosinophils and basophils (C) within *L. major* infected ear tissue at 3 wpi. Isotype controls are shown for comparison. **D-F**. Ten weeks prior to infection with *Lm*$^{DsRed}$, C57BL/6 mice were lethally irradiated and reconstituted with *Pdpn-cre* × *Il7r*$^{fl/fl}$ BM (blue line and symbols) or littermate control BM (*Pdpn-Cre* × *Il7r*$^{wt/wt}$, black line and symbols). Gating strategy for $CD4^+$ T cells in the draining lymph node (dLN) at week 3 p.i. **E.** Gating examples of infected cells in the ear tissue from mice reconstituted either with littermate control or *Pdpn-cre* × *Il7r*$^{fl/fl}$ BM. **F.** Gating strategy for effector $CD4^+$ T cells from *L. major* infected ear tissue at 3 weeks p.i. and example of IFNγ expression in either mice reconstituted with littermate control or *Pdpn-cre* × *Il7r*$^{fl/fl}$ BM. **G.** Experimental setup for *Il7r*-deficiency in non-lymphocyte immune cells. Ten weeks prior to infection with *Lm*$^{SWITCH}$, C57BL/6 mice were lethally irradiated and reconstituted with *Il7r*$^{-/-}$*Rag1*$^{-/-}$ (blue symbols) or *Il7r*$^{WT}$*Rag1*$^{-/-}$ BM (black symbols). WT $CD4^+$ T cells (pink) were adoptively transferred one day prior to infection. At 19 days p.i., *Lm*$^{SWITCH}$ were photoconverted. Applies to (H-I) **H.** Gating examples of infected cells in the ear tissue of lethally irradiated either reconstituted with *Il7r*$^{-/-}$*Rag1*$^{-/-}$ BM (upper panel) or *Il7r*$^{+/+}$*Rag1*$^{-/-}$ control (lower panel). **I.** *L. major*-infected cell fraction among monocyte-derived cells in the ear tissue (upper panel) and LDA of *L. major* tissue burden normalized to control animals (lower panel) at 3 wpi of BM reconstituted either reconstituted with *Il7r*$^{+/+}$*Rag1*$^{-/-}$ control (black symbols) or *Il7r*$^{-/-}$*Rag1*$^{-/-}$ BM (blue symbols). Each symbol represents one individual mouse ear, data pooled from two independent experiments. Horizontal lines denote the mean. ****, $p<0.0001$; * $p<0.05$ according to unpaired t-test. **J-K.** Ten weeks prior to infection with *Lm*$^{DsRed}$, C57BL/6 mice were lethally irradiated and reconstituted with either *Pdpn-cre* × iDTR BM (100% *Pdpn*-dependent depletion), 50% of *Pdpn-cre* × iDTR and 50% of CD45.1 WT BM (50% *Pdpn*-dependent depletion), or *Pdpn-cre* (iDTR-negative BM, non-depleted, also treated with DTX*)*. On 13/15/19 dpi mice were treated with DTX and analyzed 3 wpi. Applies to (D-G). **J.** Examples of the detected infected cell fractions of recruited monocyte-derived cells from ear tissue. **K.** Example of $IFN\gamma^+$ expression in $CD4^+CD44^+$ effector T cells from ear tissue from depleted (bliue line) and control (black line).

**Supplemental Figure S5: Fibroblast-produced IL-7 contributes to pathogen persistence and suppression of IFNγ production in effector T cells.**

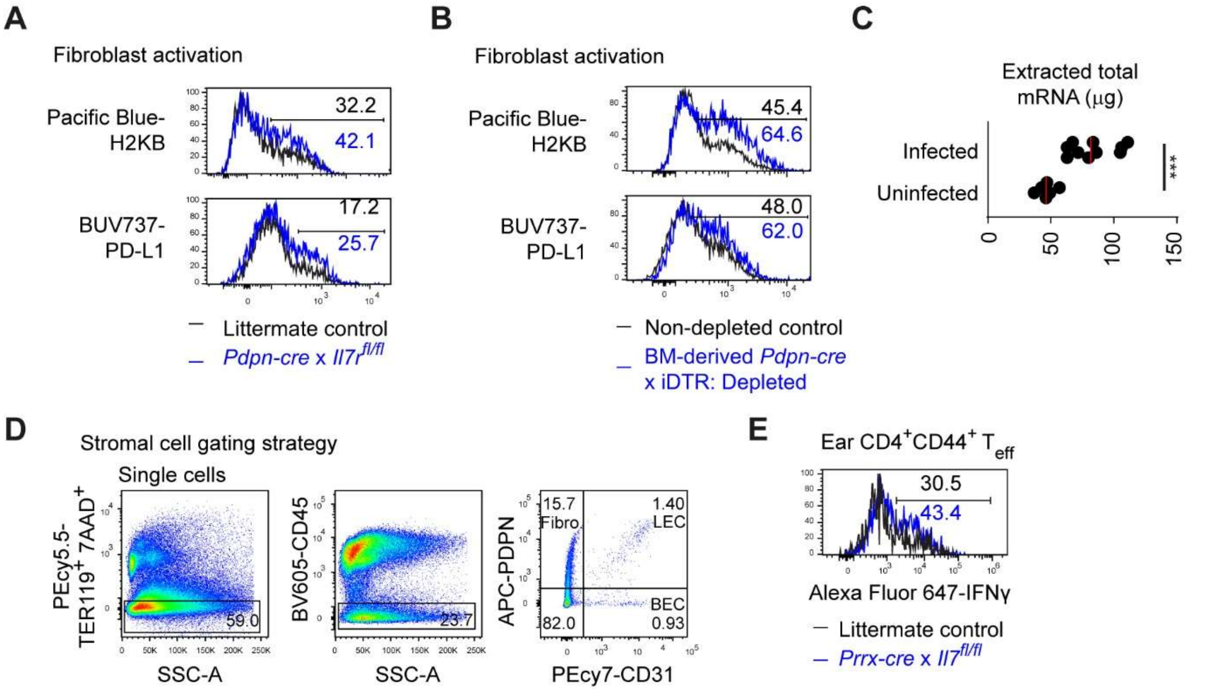


**A.** Ten weeks prior to infection with $Lm^{DsRed}$, C57BL/6 mice were lethally irradiated and reconstituted with *Pdpn-cre* × $Il7r^{fl/fl}$ BM (blue line) or *Pdpn-cre* × $Il7r^{WT/WT}$ littermate control BM (black line). $CD45^-$ $TER119^-7AAD^-PDPN^+CD31^-$ fibroblasts within infected ear tissue 3 weeks p.i. were analyzed for $H2KB^+$ (upper) and PD-$L1^+$ (lower) fractions. **B.** Ten weeks prior to infection with $Lm^{DsRed}$, C57BL/6 mice were lethally irradiated and reconstituted with *Pdpn-cre* × iDTR (blue line, 100% depleted) or *Pdpn-cre* control BM (iDTR-negative BM, non-depleted, also treated with DTX, black line). On 13/15/19 dpi mice were treated with DTX and analyzed 3 wpi (as indicated in figure 4B). $CD45^-TER119^-$ $7AAD^-PDPN^+CD31^-$ fibroblasts within infected ear tissue were analyzed for $H2KB^+$ (upper) and PD-$L1^+$ (lower) fractions. **C.** Total mRNA extracted from either *L. major*-infected (3 wpi) or uninfected C57BL/6 mouse ear tissue. Each symbol represents one individual mouse ear, data pooled from 3 independent experiments. Horizontal lines denote the median. ***, p<0.001 according to two-tailed t-test. **D.** Gating strategy for sorting fibroblasts, lymphatic endothelial cells (LEC), blood endothelial cells (BEC) and $PDPN^-CD31^-$ cells within *L. major*-infected ear tissue. **E.** Example of $IFN\gamma^+$ expression among $CD4^+CD44^+$ effector T cells within $Lm^{DsRed}$-infected ear at 3 wpi from either littermate control (*Prrx-Cre* × $Il7^{wt/wt}$; black line) or *Prrx-cre* × $Il7^{fl/fl}$ (blue line) animals.

**Supplemental Figure S6: IFNγ activates fibroblasts to produce IL-7 depending of the infection amplitude.**

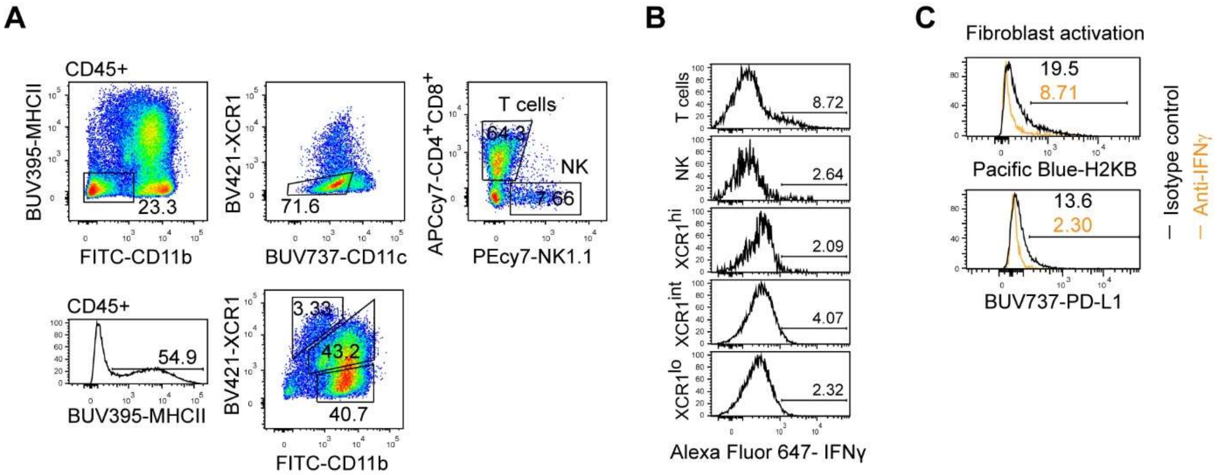


**A-B**. Gating strategy for T cells, NK cells, and XR1$^{hi/int/lo}$ antigen-presenting cells within *L. major* infected ears of C57BL/6 mice at 3 weeks p.i. (A) and examples (B) of IFNγ expression among these cell types. **C.** *Lm*$^{DsRed}$-infected C57BL/6 mice were treated with anti-IFNγ (orange lines) or isotype control (black lines) on 15 and 18 dpi**.** Example of H2KB or PD-L1 expression among fibroblasts isolated from infected ears at 3 week p.i..

**Supplemental Figure S7: Anti-IL-7/IL-7R treatment results in an enhanced immune response against *L. major*.**

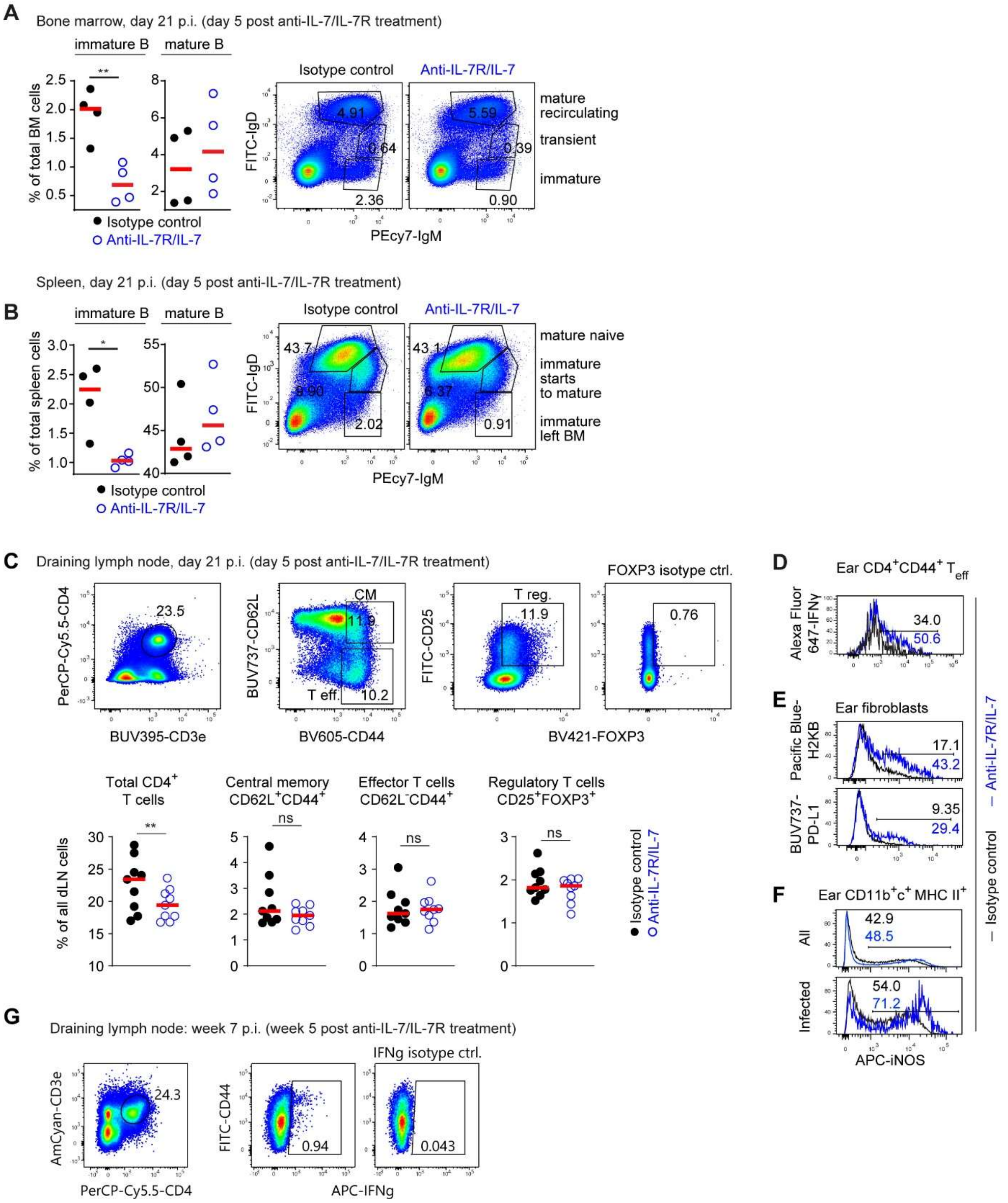


**A-B.** Fraction of immature or mature B cells among total bone marrow (**A**) or spleen (**B**) cells, respectively, following two injections of anti-IL-7R/IL-7 at 5 and 2 days before analysis. Examples of the corresponding gating strategy are shown on the right-hand side. Each symbol represents an individual mouse, data pooled from 2 independent experiments. Horizontal lines denote the mean. **,

$p<0.01$; *, $p<0.05$ according to unpaired t-test. **C.** Upper panels: Gating strategy for T cell subsets in the draining lymph node. Lower panels: Fraction of total, central memory, effector and regulatory $CD4^+$ T cells among total leukocytes in the draining dLN following two injections of anti-IL-7R/IL-7 at 5 and 2 days before analysis. Horizontal lines denote the mean. **, $p<0.01$; ns, not significant according to one-way ANOVA with Tukey's post test. Data pooled from at least 2 independent experiments. Each symbol represents one lymph node. **D.** Example of IFNγ production among effector $CD4^+$ T cells from *L. major*-infected ear tissue at 21 dpi after isotype control treatment (black lines), or anti-IL-7R/IL-7 treated (blue lines). **E.** Examples of H2KB (upper) and PD-L1 (lower) surface expression in fibroblasts within infected ear tissue after anti-IL-7R/IL-7 treatment. Blue and black lines represent the anti-IL-7R/IL-7 treated and isotype control samples, respectively. **F.** Examples of iNOS content among all recruited (upper) or infected monocyte-derived cells (lower) at 21 dpi after anti-IL-7R/IL-7 treatment. Blue and black lines represent the treatment and isotype control samples, respectively. **G.** Gating strategy for IFN$\gamma^+$ effector $CD4^+$ T cells isolated from the draining lymph node at 7 weeks p.i.
